# PAK6 promotes neuronal autophagy by regulating TFEB nuclear translocation

**DOI:** 10.1101/2024.06.05.597537

**Authors:** Francesco Agostini, Rossella Agostinis, Martina Di Rocco, Sandro Montefusco, Giulia Tombesi, Lucia Iannotta, Susanna Cogo, Federica De Lazzari, Isabella Tessari, Laura Civiero, Evy Lobbestael, Veerle Baekelandt, Gabriele Sales, Diego Luis Medina, Xiaomeng Zhang, Elizabeth Hinde, Simone Martinelli, Marco Bisaglia, Nicoletta Plotegher, Elisa Greggio

## Abstract

Autophagy is a highly conserved homeostatic process essential for the bulk degradation of cytoplasmic components and aggregated proteins. Multiple evidence indicates that impairment of (macro)autophagy leads to neurodegeneration, such as Parkinson disease (PD). Our previous work showed that p21 activated kinase 6 (PAK6) interacts with the PD-associated leucine-rich repeat kinase (LRRK2) to promote neurite outgrowth in the mouse striatum; still the function of PAK6 in the brain is largely unknown. Here, we found that downregulation of neuronal but not glial *mbt*, the *D. melanogaster* homolog of PAK6, impairs autophagy-lysosomal function. PAK6 overexpression in cells and in *C. elegans* increases transcription factor EB (TFEB) nuclear translocation in a kinase activity-dependent manner. Mechanistically, PAK6 forms a complex with TFEB to regulate its nuclear localization in a manner dependent on phosphorylation of and binding to 14-3-3 proteins and phosphorylation of TFEB at S467. In line with its ability to promote neuronal autophagy, *mbt* downregulation exacerbates alpha-synuclein toxicity in *Drosophila* dopaminergic neurons. Moreover, PAK6 overexpression in the *substantia nigra* of mutant LRRK2 mice reduces the burden of phosphorylated alpha-synuclein in dopaminergic neurons. Altogether, our study uncovers a novel role of PAK6 as a positive regulator of autophagy via TFEB and suggests that modulating its activity may represent a way to selectively turn on autophagy in neurons, with implications for the treatment of neurodegenerative disorders.

## INTRODUCTION

Macroautophagy (hereafter autophagy) is a core molecular mechanism for maintaining cellular homeostasis and for the removal of intracellular waste, such as misfolded or aggregated proteins and dysfunctional organelles [1]. The master regulator of autophagy and lysosomal biogenesis is the Transcription Factor EB (TFEB), which binds the CLEAR (Coordinated Lysosomal Expression and Regulation) gene network [2]. TFEB is regulated through posttranslational modifications, i.e. phosphorylation at serine residues, and the highly conserved pattern of TFEB phosphorylation finely controls its intracellular localization and transcriptional activity [3].

14-3-3 proteins are a family of adaptor proteins that bind phosphorylated TFEB to maintain the transcription factor in the cytoplasm. Upon calcium-mediated TFEB dephosphorylation through the phosphatase calcineurin, TFEB translocates to the nucleus and initiates its transcriptional activity [4]. The main upstream regulator of the pathway is the mechanistic target of rapamycin complex 1 (mTORC1), which phosphorylates TFEB at various serine residues, including S211, promoting the binding with 14-3-3 proteins [5]. Moreover, the mTORC1-independent phosphorylation of a stretch of serine residues localized at the C-terminus of TFEB (S466, S467 and S469) was shown to participate in the regulation of the transcription factor activity. Among the proteins that have been proposed to phosphorylate TFEB in these residues, AMP-activated protein kinase (AMPK) increases the transcriptional activity of TFEB [6].

Neurodegenerative diseases are a class of devastating disorders for which no cure is available. The majority of them, including Alzheimer disease (AD), Parkinson Disease (PD) and Amyotrophic Lateral Sclerosis (ALS), are characterized by the accumulation of protein aggregates and dysfunctional organelles accompanied by defects in the autophagic lysosomal pathway (ALP) [7]. A functional ALP is particularly crucial in non-dividing cells like neurons, which need to efficiently remove intracellular waste to maintain their homeostasis. Accordingly, knockout of key autophagy genes in mice results in neurodegeneration [8,9].

P21-activated kinases (PAK) are a family of serine/threonine kinases divided in subgroup I (PAK1-2-3) and subgroup II (PAK 4-5-6), and acting downstream of small GTPases Rac1/cdc42 [10]. Of interest, PAKs were linked to autophagy regulation in cancer [11] and more recently in neurodegeneration [10,12]. Specifically, PAK1 acetylation induced by hypoxia enhances PAK1 kinase activity, leading to phosphorylation of ATG5 that mediates autophagosome formation and promotes glioblastoma growth [11]. In contrast, PAK1 was shown to inhibit autophagy via activation of the AKT-mTOR pathway in lung and breast cancer [13,14]. Similarly, PAK2 expression was inversely correlated with autophagy induction in models of neuroblastoma [15]. A few studies linked group II PAKs with neuronal autophagy. Depletion of PAK4 in nigral dopaminergic (DA) neurons of mice causes accumulation of alpha-synuclein (α-syn), the pathological hallmark of PD, while overexpression of a constitutively active mutant of PAK4 reduces α-syn accumulation and rescues neurodegeneration in 6-hydroxydopamine and α-syn rat models for PD [16,17]. In agreement with this observation, depletion of *mbt* (mushroom bodies tiny), the *Drosophila melanogaster* (*D. melanogaster*) ortholog of group II PAKs, leads to a reduction in the number of DA neurons and a consequent impairment of the motor skills of the flies [12]. Nevertheless, these models of neurodegeneration did not provide any information on the precise effect of PAKs’ depletion or activation on autophagy.

To fill this gap, we investigated the role of group II PAKs and in particular of PAK6, in the regulation of the ALP. We used multiple cellular and animal models, including *D. melanogaster*, *Caenorhabditis elegans* (*C. elegans*) and mice, to describe a previously undisclosed role of PAK6 in regulating TFEB nuclear translocation through a mechanism involving TFEB and 14-3-3 binding and phosphorylation. In agreement with its enriched expression in neurons, the exploitation of PD models showed that depletion of *mbt* in flies led to an increase in α-syn accumulation, while overexpression of constitutively active PAK6 in the *substantia nigra* of PD mutant LRRK2-G2019S mice resulted in the reduction of the toxic phosphorylated α-syn in DA neurons. We propose PAK6 as a novel target for selectively ameliorating autophagy in neurons, as a potential strategy to counteract neurodegeneration.

## RESULTS

### Neuronal *mbt* knock-down exhibit autophagy alterations in *Drosophila* neurons

Recent studies showed that loss of *mbt*, the *Drosophila* homolog of group II PAKs, results in motor dysfunction and DA neuron degeneration that are reminiscent of PD [12]. Our previous work indicates that PAK6 kinase activity is beneficial in models of PD associated with LRRK2 mutations [18–20].

To gain more insight into group II PAKs physiological function, we downregulated *mbt* in *Drosophila* by expressing an RNAi in the whole fly body, in glial cells and in neurons, under the daughterless (da), the repo and the elav promoters, respectively (Fig. 1A). Each *Drosophila* strain was analyzed in comparison to its genetic background matched control. The selected RNAi sequences were capable of significantly downregulating *mbt* in the whole fly (Fig. S1A) and in neurons (Fig. 1B). Instead, no effect was observed in glial *mbt* knock-down (KD) flies when head lysates were analyzed by western blot. This result could be explained by the fact that glial cells represent only the 10 % of the total cell population in fruit fly brains [21].

**Figure 1.**
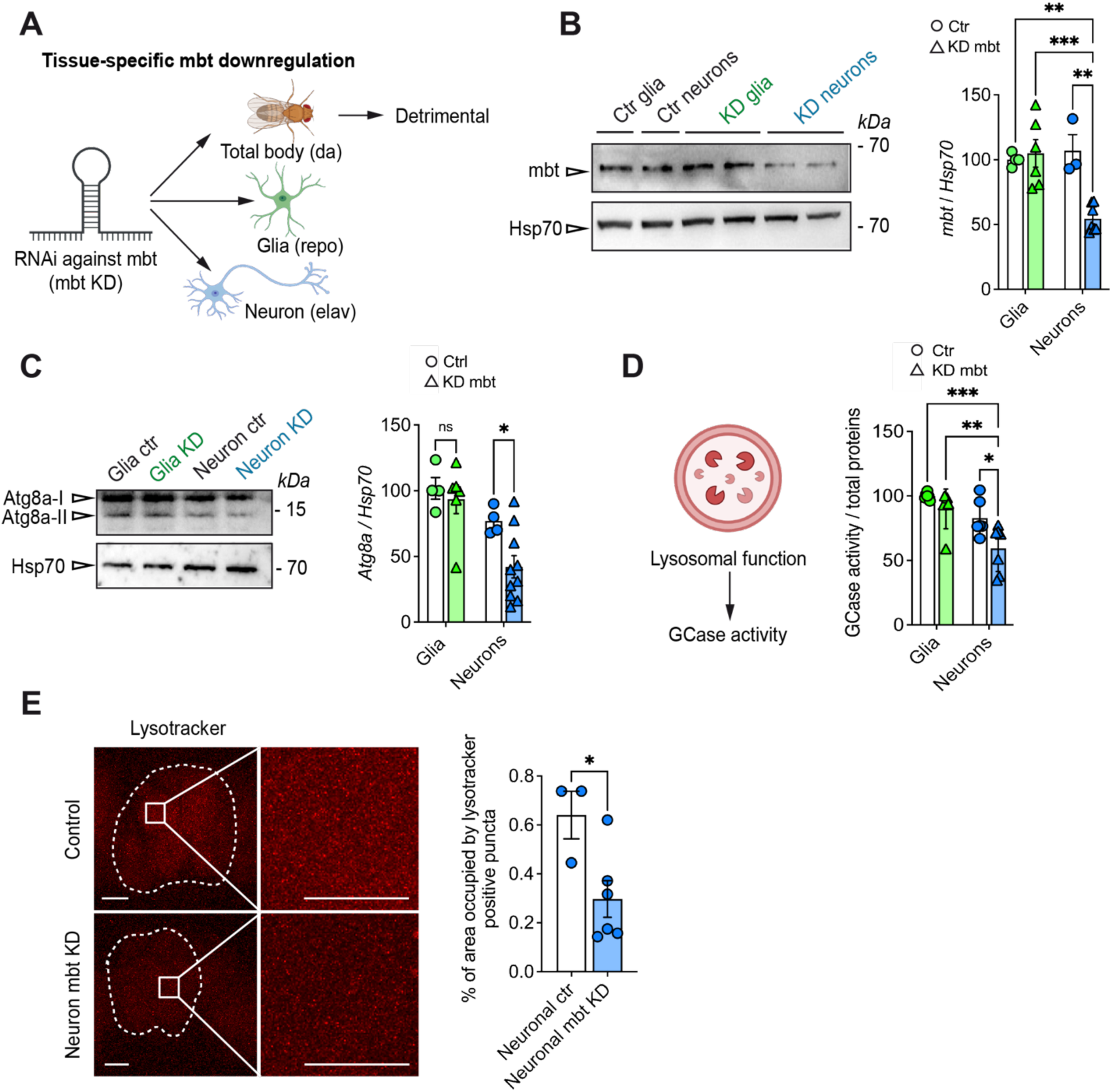
*mbt* knock-down files display autophagy alterations in neurons but not in glia cells. (**A**) Strategy of *mbt* knockdown with GAL4-UAS system expressing a RNAi against *mbt* in the total body (daughterless promoter), in neurons (elav promoter) or in the glia (repo promoter). (**B**) Western blot analysis of mbt downregulation in neurons and in glial cells. At least 3 independent samples (pool of 10 flies) were evaluated. Statistical significance was determined by two way ANOVA with Tukey’s multiple comparisons test (interaction: p=0.0036, F (1, 16) = 11.59; cell type: p=0.0203, F (1, 16) = 6.636; genotype: p=0.0120, F (1, 16) = 0.026; Glia:Ctr vs. Neurons:KD mbt ** p < 0.01; Glia:KD mbt vs. Neurons:KD mbt *** p < 0.001; Neurons:Ctr vs. Neurons:KD mbt ** p < 0.01). (**C**) Western blot analysis of Atg8a-II levels in neurons and in glial cells. At least 4 independent samples (pool of 10 flies) were evaluated. Statistical significance was determined by two way ANOVA with Šídák’s multiple comparisons test (interaction: p=0.2135, F (1, 20) = 1.651; cell type: p=0.0015, F (1, 20) = 13.53; genotype: p=0.0502, F (1, 20) = 4.342; Glia:Ctr vs. Glia:KD mbt p=0.8351; Neurons:ctr vs. Neurons:KD mbt *p < 0.05). (**D**) GCase enzymatic assay of fly heads downregulating mbt in neurons vs. glial cells and relative matching controls. N=6-7 biological replicates. Statistical significance was determined by two way ANOVA with Tukey’s multiple comparisons test (interaction: p=0.2424, F (1, 21) = 1.447; cell type: p=0.0004, F (1, 21) = 18.13; genotype: p=0.0071, F (1, 21) = 8.903; Glia:Ctr vs. Neurons:KD mbt ***p < 0.001; Glia:KD mbt vs. Neurons:KD mbt **p < 0.01; Neurons:Ctr vs. Neurons:KD mbt *p < 0.05). (**E**) Lysotracker red of neuronal mbt KD brains (scale bar 50 μm, scale bar magnification 20 μm). The area occupied by lysotracker-positive structures was quantified. Statistical significance was determined by unpaired t-test (*p>0.05) using at least 3 brains per genotype.

As shown by the lifespan experiments performed on the three different *mbt* KD lines (Fig. S1B-C-D), whole-body *mbt* KD led to a ~50% reduction in the survival rate after only 20 days (Fig. S1B). The whole-body KD of *mbt* also impacted the overall fly development, causing severe climbing defects, as measured by the negative geotaxis assay (Fig. S1E) and decreased hatching (Fig. S1F). Taken together, these data confirm previous studies showing robust developmental and phenotypic defects in flies downregulating *mbt* in the whole body [12]. Thus, the subsequent investigations were performed in brain cell-type specific *mbt* KD flies. Firstly, we quantified the levels of Atg8a (LC3) in the heads of neuronal and glial *mbt* KD flies and the activity of the enzyme glucocerebrosidase (GCase) as readout of lysosomal function. As shown in figure 1C, *mbt* KD causes a reduction in the lipidated form of Atg8a in neurons but not in glial cells, and parallel decrease in the GCase activity, which were observed again only in *mbt* KD neurons (Fig. 1D), suggesting defective lysosomal activity or a reduced lysosomal compartment. To further dissect these two possibilities, neuronal *mbt* KD brains were stained with lysotracker and the amount of lysosomes evaluated by confocal microscopy imaging. As figure 1E illustrates, the area occupied by lysotracker staining was reduced in *mbt* KD brains compared to controls, demonstrating a decreased number of lysosomes upon *mbt* downregulation. Taken together these data support an involvement of *mbt* in the regulation of ALP in *Drosophila* neurons.

### PAK5 and PAK6 are highly expressed in the brain

*Drosophila mbt* underwent gene duplication resulting in three paralogs in mammals [22]. Based on the autophagy-lysosome phenotype observed in neuron-specific *mbt* KD flies (Fig. 1), we next analyzed the co-expression patterns of group II PAKs (PAK4, PAK5 and PAK6) using the Human Protein Atlas consensus data, which integrates gene expression from 50 healthy tissues (Fig. 2A). As shown in the heatmaps of the top 50 co-expressed genes, PAK4 is ubiquitously expressed, while PAK5 and PAK6 are highly enriched in brain cells/tissues (Fig. 2A, Fig. S2A-C). Next, we subjected the top 50 co-expressed genes to functional enrichment analysis with gProfiler (g:GOSt). Gene ontology identified only one term for PAK4 (GO:BP = tRNA-type intron splice site recognition and cleavage with *p* = 0.024517272), confirming the poor tissue specificity of this member [23]. Strikingly, PAK5 and PAK6 co-expressed genes are instead enriched for biological processes related to synaptic structure/function, strongly supporting a synapse-specific function of the two kinases (Fig. 2A). To increase the specificity, we selected only those GO categories with a term size < 500 and the top 8 GO:BP categories were plotted (Fig. 2B). Given the selective enrichment for synaptic-related BP terms, we then subjected the 50 co-expressed genes to SynGO [24] and found that 26/50 PAK5 and 14/50 PAK6 co-expressed genes are annotated in SynGO, falling into both pre- and post-synaptic categories (Fig. 2C). Altogether, these analyses indicate that PAK5 and PAK6 are neuronal kinases enriched at the synapses.

**Figure 2.**
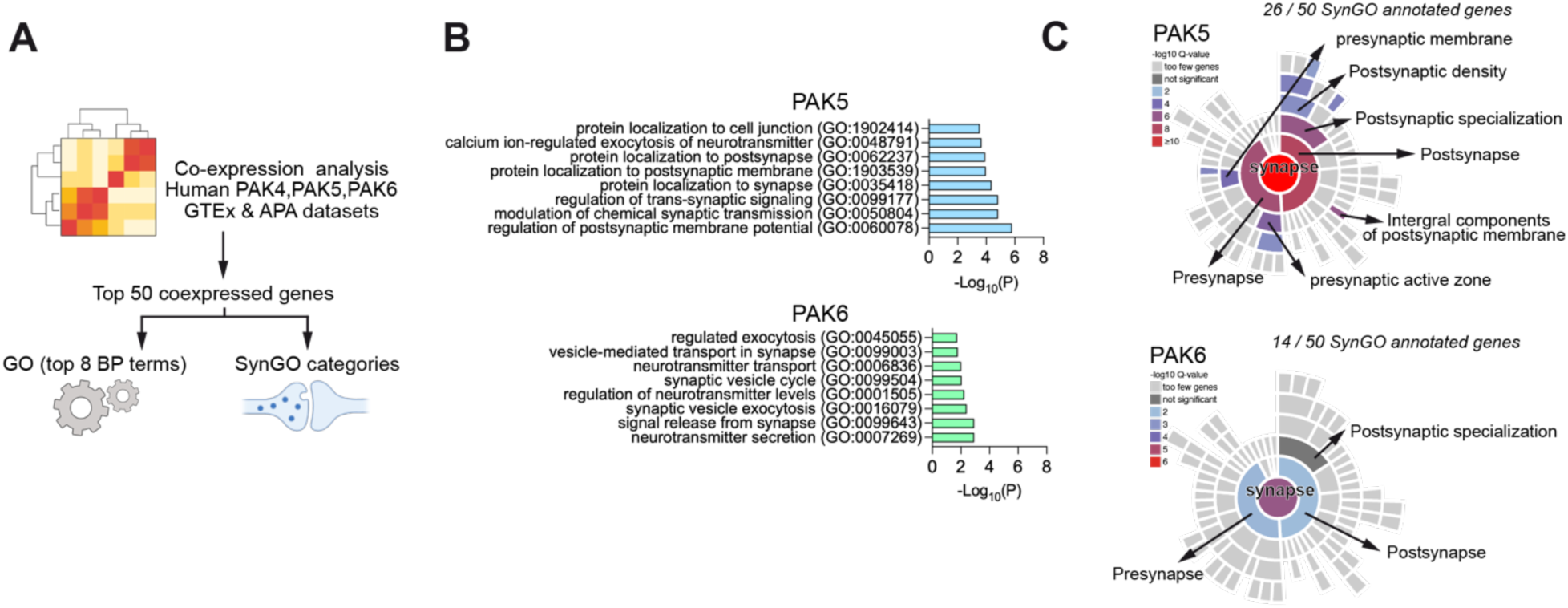
PAK5 and PAK6 are highly expressed in the brain. (**A**) Schematic representation of co-expression and gene ontology analyses. (**B**) G:profiler g:GOSt pathway enrichment analysis showing the top 8 enriched categories of PAK5 and PAK6 co-expressed genes. (**C**) Sunburst plot showing enriched SynGO (https://www.syngoportal.org/) categories in PAK5 (top) and PAK6 (bottom) top 50 co-expressed genes.

### PAK6 responds to mTORC1 inhibition and increases TFEB levels and nuclear compartmentalization in neuronal cells

Since PAK6 has been previously linked to LRRK2-related PD [18–20] and its expression is predicted to be influenced upon TFEB overexpression, while PAK5 (also known as PAK7) is not affected (https://tfeb.tigem.it/index.php, citation:[25]), we focused on PAK6 for the subsequent mechanistic studies. We first investigated whether the kinase is involved in the canonical autophagic pathway regulated by mTORC1. To this end, we assessed the impact of mTORC1 inhibition on PAK6 activation, by evaluating the level of autophosphorylated (active) form of the kinase through confocal imaging (Fig. 3A) and immunoblot analysis (Fig. 3B) in neuroblastoma SH-SY5Y cells untreated or treated with the specific mTORC1 inhibitor and strong autophagy inducer Torin1. Upon mTORC1 inhibition, we detected a significant enhancement in PAK6 activity, as indicated by the increase in the phosphorylation of Ser560, both in naïve and in PAK6 stably overexpressing (OE) cells. The phospho-antibody used is not specific for PAK6, as this phospho-site is conserved in PAK4, PAK5 and PAK6; nevertheless, the use of cells overexpressing PAK6 and the significant difference in the antibody signal between the two lines indicate that the observed effects are PAK6-dependent. These results strongly support PAK6 as a downstream component of mTORC1 pathway.

**Figure 3.**
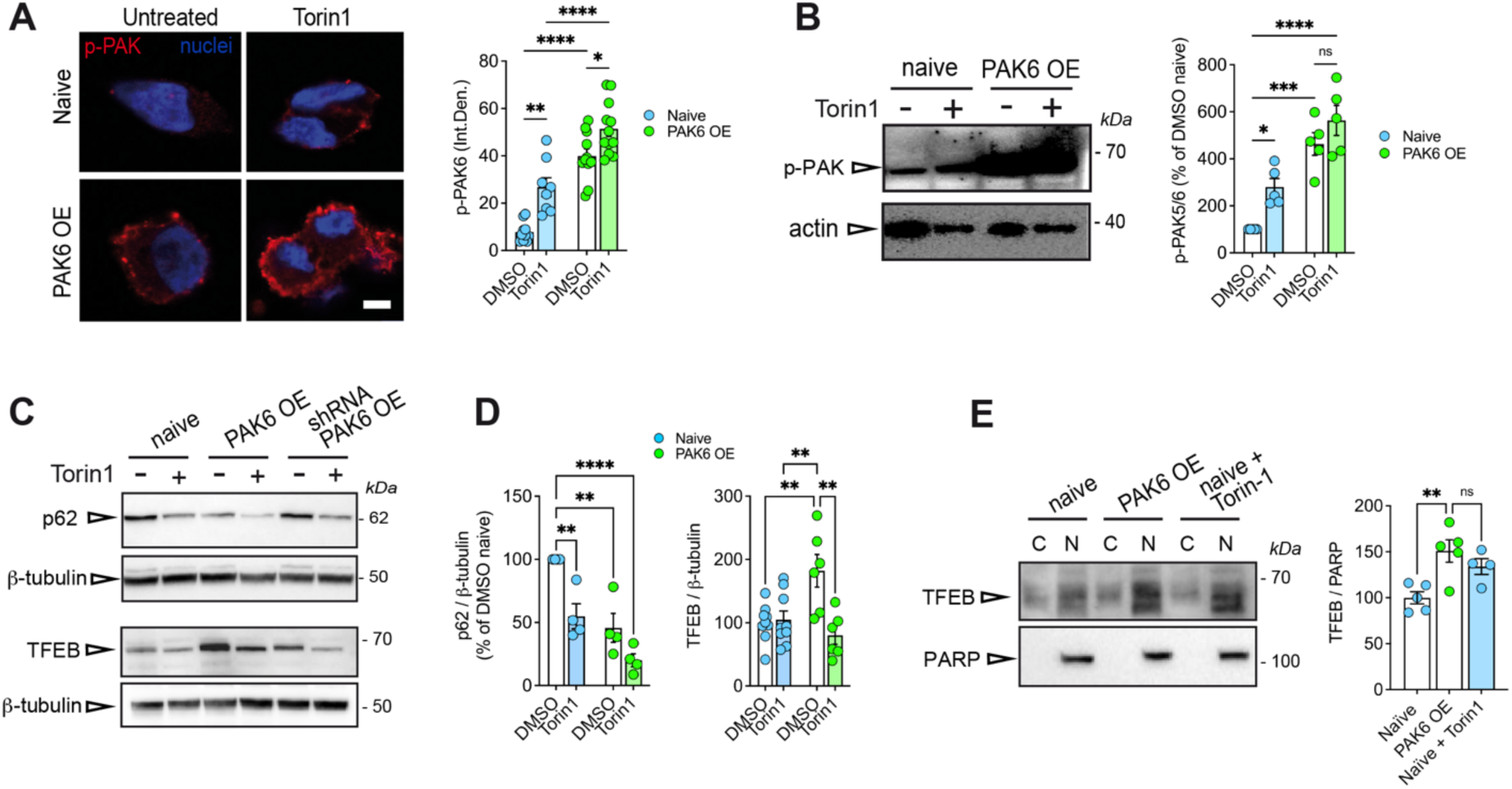
PAK6 responds to mTORC1 inhibition and increases TFEB levels and nuclear localization in neuronal cells. (**A**) Confocal imaging of PAK6 autophosphorylation at Ser560 upon inhibition of mTORC1 via Torin1 treatment (2.5 μM for 90 minutes) in SH-SY5Y naïve or stably overexpressing PAK6 (PAK6 OE) and confocal imaging (scale bar: 10 μm). Right: quantification of p-PAK signal (integrated density) of 8-12 cells per experiment. Statistical significance was determined by two-way ANOVA with Tukey’s multiple comparisons test (interaction: p=0.2317, F (1, 37) = 1.479; treatment p<0.0001, F (1, 37) = 24.98; genotype: p<0.0001, F (1, 37) = 86.63; DMSO:naïve vs. DMSO:PAK6 OE ****p<0.0001; DMSO:naïve vs. Torin1:naïve **p<0.01; DMSO:naïve vs. Torin1:PAK6 OE ****p<0.0001; DMSO:PAK6 OE vs. Torin1:naïve *p<0.05; DMSO:PAK6 OE vs. Torin1:PAK6 OE *p<0.05; Torin1:naïve vs. Torin1:PAK6 OE ****p<0.0001). (**B**) Western blot showing PAK6 autophosphorylation at Ser560 (anti-PAK4/5/6 antibody) upon inhibition of mTORC1 via torin1 treatment (2.5 μM for 90 minutes) in SH-SY5Y naïve or stably overexpressing PAK6. Right: quantification of n=5 replicates. Statistical significance was determined by two-way ANOVA with Tukey’s multiple comparisons test (interaction: p=0.3773, F (1, 16) = 0.8247; treatment: p=0.0057, F (1, 16) = 10.20; genotype: p<0.0001, F (1, 16) = 54.11; DMSO:naïve vs. DMSO:PAK6 OE ***p<0.001; DMSO:naïve vs. Torin1:naïve *p<0.05; DMSO:naïve vs. Torin1:PAK6 OE ****p<0.0001; DMSO:PAK6 OE vs. Torin1:naïve *p<0.05; Torin1:naïve vs. Torin1:PAK6 OE **p<0.01). (**C**) P62 and TFEB levels in SH-SY5Y cells (naïve, PAK6 overexpressing OE, shRNA-PAK6 OE) stimulated or not with torin1 (2.5 μM for 90 minutes). (**D**) Quantification of p62 and TFEB levels in SH-SY5Y cells (naïve and PAK6 OE) stimulated or not with torin1 (2.5 μM for 90 minutes). Statistical significance was determined by two-way ANOVA with Tukey’s multiple comparisons test (TFEB levels: interaction p=0.0022, F (1, 26) = 11.52; treatment p=0.0048, F (1, 26) = 9.510; genotype p=0.0800 F (1, 26) = 3.320. P62 levels: interaction p=0.2477, F (1, 12) = 1.476; treatment p=0.0008, F (1, 12) = 19.59; genotype p=0.0001, F (1, 12) = 31.21. Multiple comparisons: *p<0.05, **p<0.01; ***p<0.001; ****p<0.0001). (**E**) Western blot of nuclear vs. cytosolic fractions from SH-SY5Y cells naïve, PAK6 OE or naïve treated with Torin-1 (2.5 μM for 90 minutes) probed with anti-TFEB and anti-PARP (nuclear marker) antibodies. Right: quantification of the ratio between nuclear TFEB and PARP (n=4-5). One-way ANOVA with Tukey’s multiple comparisons test (**p<0.01).

To further characterize the contribution of PAK6 in the regulation of autophagy, we evaluated the levels of p62 in naïve and PAK6 OE cells. Interestingly, we detected a decreased steady state level of p62 in PAK6 OE cells, suggesting that PAK6 activity may promote the autophagic degradation of this protein. The effect was comparable to the one obtained upon treatment with the autophagy inducer Torin1 (Fig. 3C-D). In addition, we observed that the overexpression of PAK6 resulted in a two-fold increase in TFEB protein levels, pointing to PAK6 as a potential regulator of the transcriptional induction of autophagy (Fig. 3 C-D). Importantly, we were able to restore the control phenotype by downregulating PAK6 in PAK6 OE cells, linking the observed results to the activity of PAK6 (Fig. 3C). When activated, TFEB enters the nucleus to promote the transcription of the ALP-related genes [26]. To evaluate whether PAK6 also promotes TFEB nuclear translocation, we performed nuclear fractionation and TFEB immunoblotting in naïve (+/- Torin1) and PAK6 OE cells. As shown in figure 3E, the nuclear amount of TFEB positively correlates with the level of PAK6, supporting a mechanism whereby PAK6 promotes TFEB nuclear translocation.

### PAK6 kinase activity promotes TFEB nuclear translocation *in vitro* and *in vivo*

To further explore the impact of PAK6 kinase activity on TFEB nuclear translocation, we performed confocal imaging in the presence of overexpressed PAK6 wild-type (WT), constitutively active (S531N) or kinase dead (K436M) alongside GFP-TFEB and correlated the level of PAK6 activity with the level of TFEB in the nucleus. Constitutively active PAK6^S531N^ dramatically increased the amount of nuclear TFEB, comparable in magnitude to the effect observed with Torin1 treatment (Fig. S3A-B). Conversely, exogenous expression of PAK6^K436M^ completely prevents the shuttling of the transcription factor from the cytoplasm to the nucleus, which is comparable to the rate of TFEB nuclear translocation observed in cells transfected with a control plasmid (Gus) (Fig. S3A-C). However, since PAK6 WT did not result in any clear effect on overexpressed GFP-TFEB, we developed a high-content imaging assay to measure PAK6-dependent TFEB nuclear translocation under physiological levels of TFEB in HeLa cells. PAK6 WT and, at a greater extent PAK6^S531N^, significantly increased endogenous TFEB nuclear translocation as compared to inactive PAK6^K436M^ or mCherry control (Fig. 4A-B). Altogether these results support PAK6 activity as a novel regulator of TFEB nuclear translocation *in vitro*.

**Figure 4.**
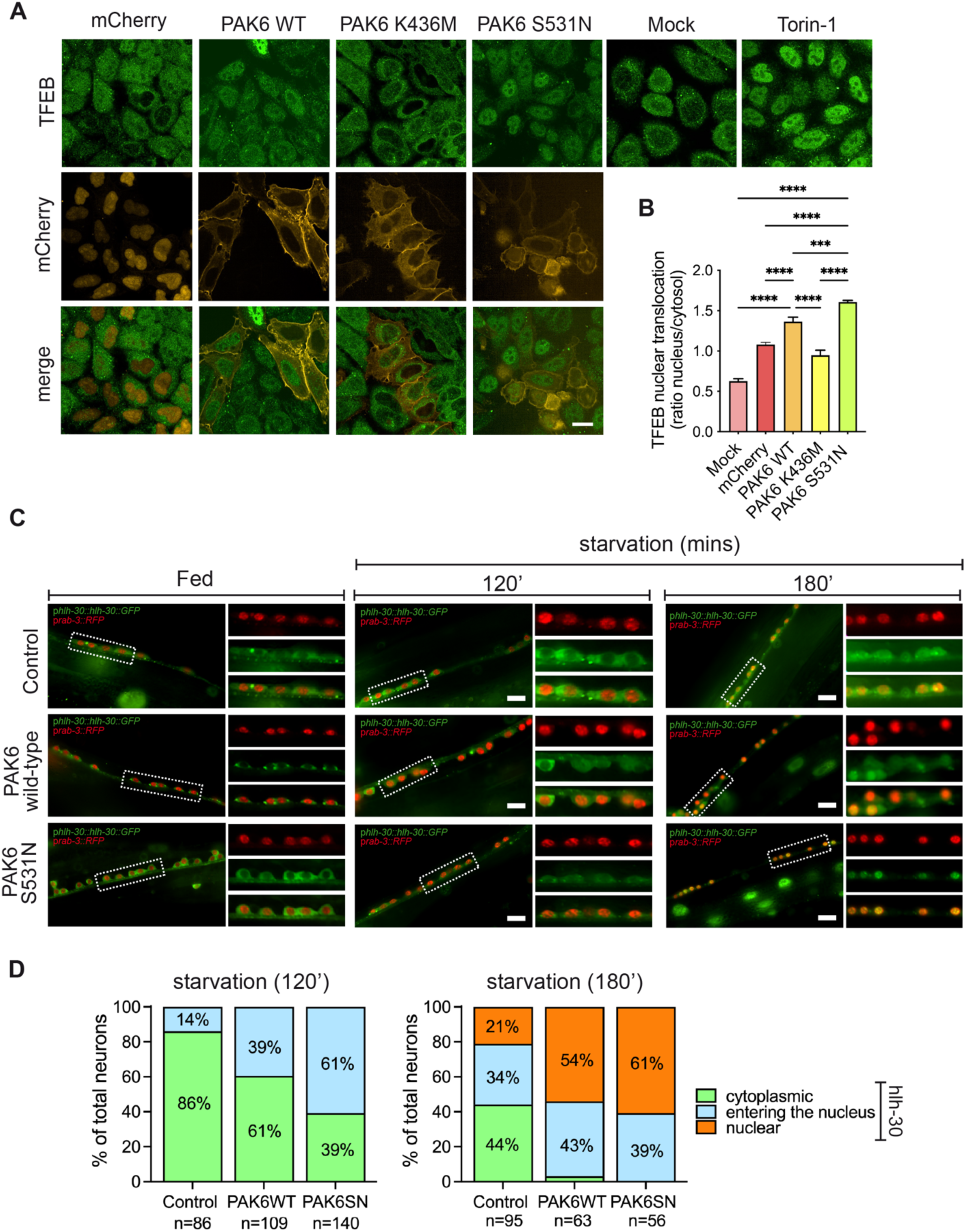
PAK6 kinase activity induces TFEB nuclear translocation *in vitro* and *in vivo.* (**A**) Endogenous TFEB nuclear translocation in HeLa cells transfected with mCherry, PAK6 WT, PAK6 S531N, and PAK6 K431M (Scale bar 10 μm). (**B**) Quantification of n=3 independent experiments. Number of transfected cells analyzed per each experiment: mCherry (577, 552, 413); PAK6-WT (348, 468, 309); PAK6-K436M (166, 232, 229); PAK6-S531N (136, 198, 226). Statistical significance was determined using one-way ANOVA with Turkey’s multiple comparisons test (****p<0.0001). (**C**) PAK6 induces HLH-30/TFEB nuclear translocation *in vivo*. Ectopic expression of hPAK6 WT in animals expressing GFP-tagged HLH-30/TFEB promotes translocation of HLH-30 into the nucleus (RFP-tagged) of ventral cord motor neurons after short-term starvation (120 and 180 minutes), whereas it has no effect in well-fed animals. A significantly greater effect was observed in worms expressing constitutively active hPAK6 (S531N). Scale bars 10 µm. (**D**) Bar graphs showing the percentage of neurons with HLH-30 localized in the cytoplasm, entering into the nucleus, or localized in the nucleus, in fed animals as well as after 120 or 180 minutes of starvation. Differences are quantified using Fisher’s exact test with Bonferroni correction for multiple comparisons. Specifically, after 120 minutes starvation (cytoplasmic *vs*. entering into the nucleus): WT *vs*. control p<0.005; S531N *vs*. control p<0.0001; WT *vs*. S531N p<0.005. After 180 minutes starvation (cytoplasmic *vs*. others): WT *vs*. control p<0.0001; S531N *vs*. control p<0.0001; WT *vs*. S531N non-significant. After 180 minutes starvation (cytoplasmic *vs*. nuclear): WT *vs*. control p<0.0001; S531N *vs*. control p<0.0001; WT *vs*. S531N non-significant. After 180 minutes starvation (entering into the nucleus *vs*. nuclear): WT *vs*. control non-significant; S531N *vs*. control p<0.05; WT *vs*. S531N non-significant. Ten animals and 90 neurons for each condition and genotype were tested on average.

To validate the capacity of PAK6 to promote TFEB nuclear translocation *in vivo*, we employed *C. elegans* as a model system. In the nematode, HLH-30, the ortholog of mammalian TFEB, is a master regulator of autophagy-related genes belonging to the CLEAR network [27]. HLH-30 also controls resistance to stressors, including starvation, heat shock, oxidative stress, and pathogen infection [28,29], and modulates longevity [27]. Sequence alignment highlighted conservation of several key residues mediating cellular function between human and worm proteins (e.g. Ser134, Ser211, Ser459, Ser463, Ser466, Ser469) (Fig. S4A). As expected, HLH-30 localized to the cytoplasm in well-fed animals (Fig. 4C) but translocated into the nucleus in the absence of food (overnight starvation) (Fig. S4B). To explore the role of PAK6 in HLH-30 activation, we generated transgenic lines overexpressing the wild-type human PAK6 (hPAK6) protein or the hyperactive S531N mutant in *C. elegans* neurons. Fluorescent microscopy analysis did not show any effect of ectopic hPAK6 expression in fed animals (100% cytoplasmic in controls and both transgenic lines) (Fig. 4C). In contrast, following short-term starvation (120 and 180 minutes), overexpression of hPAK6 WT promoted translocation of HLH-30 into the nucleus of ventral cord motor neurons (Fig. 4C-D). Of note, a stronger effect was observed in worms overexpressing constitutively active hPAK6^S531N^. In both transgenic lines, the most robust effect was observed after a 3-hour starvation. These findings support the role of PAK6 in facilitating TFEB nuclear translocation *in vivo*. Moreover, since the expression of human PAK6 influences the behaviour of the worm TFEB orthologue, these data also highlight a conserved function of PAK6 across evolution.

### PAK6 forms a complex with TFEB to regulate its nuclear import in a manner dependent on phosphorylation of and binding to 14-3-3 proteins and phosphorylation of TFEB at S467

TFEB nuclear translocation is triggered by calcineurin-dependent dephosphorylation of TFEB and consequent dissociation of 14-3-3 proteins [4]. PAK6 phosphorylates 14-3-3s at S58/59 and 14-3-3s bind PAK6 at S113 autophosphorylation site (Fig. 5A;[19]). To explore the mechanism whereby PAK6 affects TFEB nuclear transport, we initially evaluated whether PAK6 activity could influence TFEB phosphorylation by monitoring the electrophoretic mobility of phospho-TFEB, which is known to be slower compared to the unphosphorylated form [30]. Under unstimulated conditions, both naïve and PAK6 OE cells showed a fraction of phosphorylated TFEB in western blot (upper band, Fig. 5B), which was abolished in naïve cells upon Torin1 treatment, as expected (Fig. 5B-C). Instead, the effect of Torin1 in reducing phospho-TFEB in PAK6 OE cells was significantly smaller (*P* <0.05 naïve+Torin1 vs. PAK6 OE+Torin1), suggesting that a fraction of phospho-TFEB is insensitive to mTORC1 inhibition when PAK6 is present. Thus, to test the possibility that PAK6 phosphorylates TFEB, we analyzed the presence of candidate PAK6 consensus site(s) within TFEB. Notably, we found that S467 is localized in a PAKs consensus motif and is included in an evolutionary conserved stretch of serine residues, which is also present in *C. elegans* (Fig. 5D and Fig. S4A). To investigate the putative contribution of S467 phosphorylation on PAK6-mediated regulation of TFEB, we performed the well-established GFP-TFEB nuclear translocation assay comparing WT *vs*. phospho-deficient S467A in the presence or absence of active PAK6. Consistent with PAK6 phosphorylating (or indirectly controlling the phosphorylation of) S467, the fraction of nuclear TFEB^S467A^ was lower compared to the one of TFEB WT upon PAK6^S531N^ overexpression (Fig. 5E-F). However, in the absence of PAK6 there was more TFEB^S467A^ than TFEB WT in the nucleus, suggesting the possibility that TFEB and PAK6 also form a physical complex (Fig. 5F).

**Figure 5.**
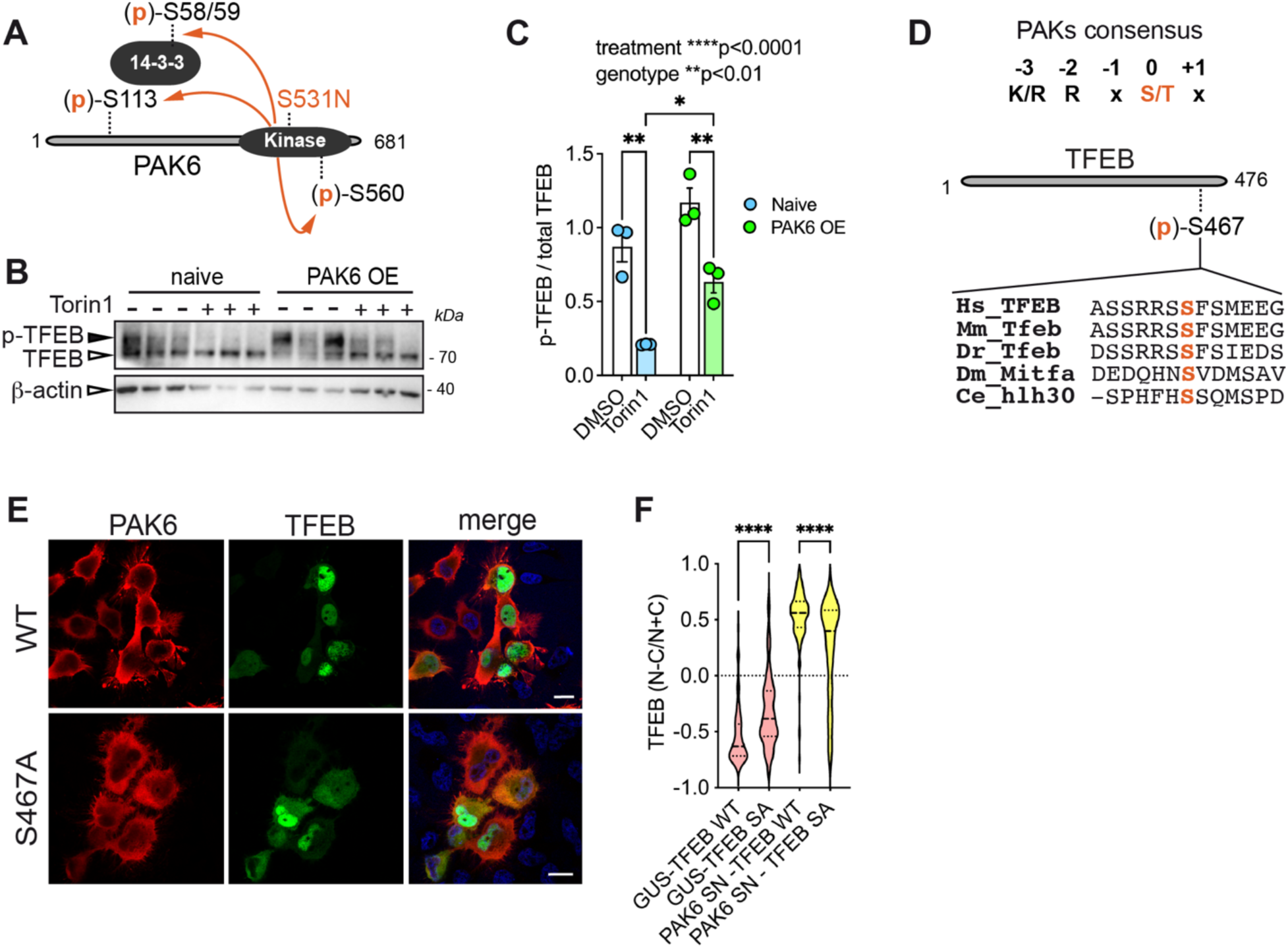
PAK6 kinase activity, binding to 14-3-3 and phosphorylation of TFEB-S467 are all positively correlated with PAK6-dependent TFEB nuclear translocation. (**A**) Schematic of PAK6-mediated phosphorylation events. S531N is a substitution that produces a constitutively active kinase [48]; S560 and S113 are two autophosphorylation sites [19]; phospho-S113 is a 14-3-3 binding site [19]. (**C**) Western blot of naïve and PAK6 overexpressing SH-SY5Y cells treated with Torin1 or vehicle control for 90 minutes. The upper band (solid arrowhead) represents phosphorylated TFEB [49]. Quantification of N=3 independent replicates per group. Statistical significance was determined by two-way ANOVA with Tukey’s multiple comparisons test (interaction p=0.4506, F (1, 8) = 0.6289; treatment ****P<0.0001, F (1, 8) = 55.99; genotype **P=0.0020, F (1, 8) = 20.38). (**D**) Schematic of PAKs consensus site and aminoacid sequence around TFEB S467, which is evolutionary conserved in the models used across this study. (**E**) TFEB nuclear translocation assessed in Hela cells cotransfected with GFP-TFEB WT or S467A (SA) and Flag-Gus or PAK6 S531N (SN) (Scale bar 10 μm). On the right quantification of Gus-TFEB WT=134; Gus-TFEB SA=150; PAK6 SN-TFEB WT=124; PAK6 SN-TFEB WT=396; Statistical significance was determined using one-way ANOVA with Tukey’s multiple comparisons test (****p<0.0001).

### Fluorescence correlation imaging in live cells supports the formation of a complex between TFEB and active PAK6

To gain a mechanistic understanding of TFEB cytosol-nuclear dynamics in the presence of PAK6, we coupled methods based on fluorescence correlation spectroscopy (FCS) with live cell imaging (Fig. S5). In particular, we acquired dual-channel line scan FCS measurements across the nuclear envelope of HeLa cells co-expressing GFP-TFEB WT or GFP-TFEB^S467A^ alongside mCherry-tagged PAK6 WT (PAK6-mCh), PAK6^S531N^ (constitutively active) (PAK6^S531N^-mCh) or PAK6^S531N/S113A^ (constitutively active and 14-3-3 deficient binding) (Fig. 6A). Then, for each condition, we calculated the autocorrelation function (ACF) in the GFP and mCherry channels (green and red plots), versus the cross-correlation function (CCF) between these two channels (yellow plots), across the cytosol versus nuclear compartment (Fig. 6B); and in those case where a complex was formed the fraction of molecules forming a TFEB-PAK6 complex (CC) was quantified (Fig. 6C) as well as their resulting diffusion properties (Fig. S6). Then finally, we also calculated the pair correlation function (pCF) versus cross pair correlation function (cross pCF), in an analogous manner to the ACF and CCF, but across the nuclear envelope in the cytoplasm-to-nucleus versus nucleus-to-cytoplasm direction, and in each case quantified the distribution of arrival times for each TFEB-PAK6 complex to perform this long-range diffusive transit (Fig. S7).

**Figure 6.**
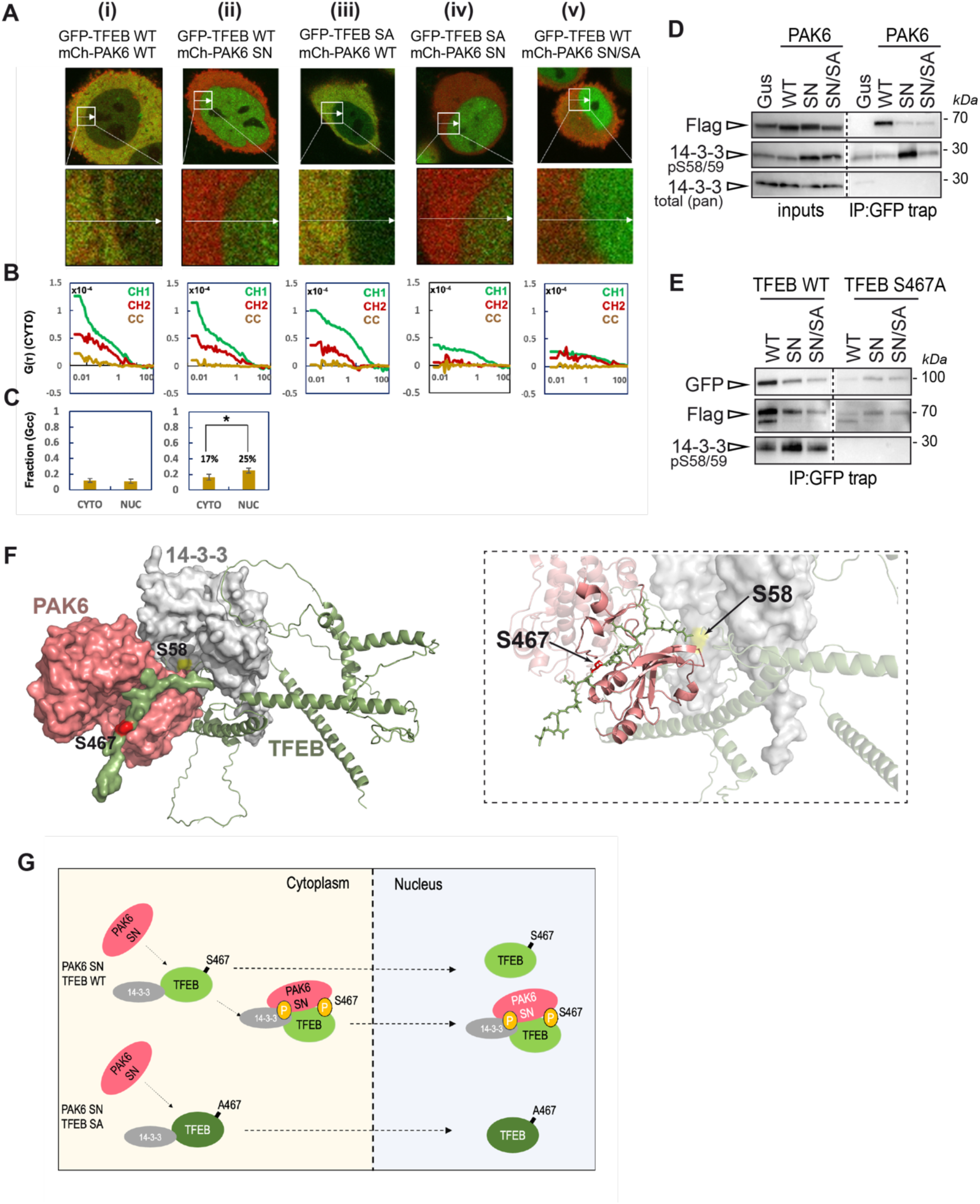
Fluorescence correlation analysis of TFEB-PAK6 nucleocytoplasmic transport. (**A**) Two channel intensity images of HeLa cells co-expressing GFP-TFEB in the presence of mCh-PAK6, mCh-PAK6-SN or mCh-PAK6-KM versus GFP-TFEB-SA in the presence of mCh-PAK6 or mCh-PAK6-SN (top row) alongside the region of interest (ROI) selected for a line scan acquisition that records transport with respect to the nuclear envelope (bottom row). (**B**) Representative G(ρ) auto correlation (channel 1 green TFEB and channel 2 red PAK6) and cross correlation functions (CC yellow) calculated from the line scan acquisitions presented in panel (A) that were used to calculate the fraction of each type of TFEB-PAK6 complex in the cytoplasm versus nucleus. (**C**) Fraction of each TFEB-PAK6 complex formed in the cytoplasm versus nucleus across multiple HeLa cells (N = 3-8 cells). (**D**) GFP-trap co-immunoprecipitation of GFP-TFEB co-expressed with Gus control, PAK6 WT, PAK6 S531N (SN), PAK6 S531N/S113A (SN/SA) in HEK293T cells. Only PAK6 WT but not PAK6 SN nor PAK6 SN/SA co-precipitates with TFEB. Both PAK6 SN and PAK6 SN/SA phosphorylate endogenous 14-3-3s at S58/S59 but only PAK6 SN and not SN/SA promotes phospho-14-3-3 co-precipitation with GFP-TFEB. Western blot were performed with anti-flag (Gus or PAK6), anti-phospho-S58/59 (14-3-3s) and anti-14-3-3s antibodies. (**E**) GFP-trap co-immunoprecipitation of GFP-TFEB WT or GFP-TFEB S467A (SA) co-expressed with Gus control, PAK6 WT, PAK6 S531N (SN), PAK6 S531N/S113A (SN/SA) in HEK293T cells. Only TFEB WT but not TFEB SA co-precipitates with phosphorylated 14-3-3s. Western blot were performed with anti-flag (Gus or PAK6), anti-phospho-S58/59 (14-3-3s) and GFP (TEFB) antibodies. (**F**) Alpha-fold2 three-component complex of TFEB (green), PAK6 (red) and 14-3-3 (grey). Residues S467-TFEB and S58-14-3-3 are highlighted. (**G**) Proposed mechanism of PAK6-dependent TFEB nuclear translocation.

As can be seen from the resulting ACF and CCF analysis (Fig. 6C): (1) PAK6 WT forms a complex with TFEB in both the cytoplasm and nucleus, while constitutively active PAK6^S531N^ promotes significantly more complex formation with TFEB in the nucleus than in the cytoplasm, and PAK6^S531N/S113A^ inhibits complex formation with TFEB throughout both compartments; (2) neither PAK6 WT or PAK6^S531N^ show complex formation with TFEB^S467A^; thus active PAK6 needs to bind phospho-TFEB to form a complex. pCF and cross pCF analysis (Fig. S7) further indicate that: PAK6 WT in complex with TFEB WT is only exported from the nucleus, while PAK6^S531N^ in complex with TFEB WT is preferentially imported into the nucleus. Interestingly, diffusion coefficients were calculated by fitting the autocorrelation curves, providing an estimate of the mode of diffusion of TFEB and PAK6. GFP-TFEB diffuses with a 2-component model, while PAK6 diffuses with a 1-component model. This suggests that TFEB translocation to the nucleus occurs in two different ways, and that one or both are at least in part regulated by PAK6 kinase activity (Fig. S5).

14-3-3 proteins are crucial regulators of TFEB subcellular localization [5] and are established substrates of PAK6 [19] (Fig. 5A). To clarify why active PAK6^S531N^ is capable of forming a complex with TFEB WT but not with TFEB^S467A^ (Fig. 6B-C), but still promoting translocation of TFEB ^S467A^ although at lesser extent (Fig. 5E-F), we hypothesized a possible involvement of 14-3-3 proteins in the PAK6-mediated regulation of TFEB. To this aim, we carried out a TFEB pulldown assay in cells co-transfected with GFP-TFEB WT together and PAK6 WT, PAK6^S531N^ or the double PAK6^S531N/S113A^ mutant, which maintains phosphorylation activity but is unable to bind 14-3-3s (Fig. 6E and [19]). GFP-trap pulldowns showed that PAK6 co-precipitates with TFEB. Noteworthy, the interaction is stronger with PAK6 WT compared to PAK6^S531N^ mutants, suggesting that increased PAK6 activity may cause a partial dissociation of the kinase from the transcription factor. Importantly, when we checked for the presence of 14-3-3s associated to TFEB, we found a dramatic increase in the binding between the transcription factor and phospho-S58/59-14-3-3s in the presence of PAK6^S531N^, but not with the 14-3-3-deficient binding mutant PAK6^S531N/S113A^ (Fig. 6E). This result is also supported by the reduced cross-correlation function observed in the presence of the 14-3-3-deficient binding mutant PAK6^S531N/S113A^ (Fig. 6Av-Cv). Altogether, these data support a mechanism whereby the active state of PAK6 and its ability to bind 14-3-3s synergize to promote TFEB nuclear localization. Given that active PAK6^S531N^ facilitates the translocation of TFEB from the cytoplasm to the nucleus (Fig. 6A-D) and phospho-14-3-3 copurifies with TFEB WT in the presence of PAK6^S531N^ (Fig. 6D-E), one possibility is that the phosphorylation of 14-3-3s at S58/59 by PAK6 triggers the formation of a three-component complex.

To explore this possibility, we employed a computational approach to model a putative tripartite complex among phospho-TFEB, phospho-14-3-3 and PAK6. We downloaded the PAK6 and 14-3-3 models from the Swiss-model database [31], and we applied the AlphaFold 2 computational method to predict the folding of TFEB. We then used the ClusPro web service [32] to evaluate the docking of these proteins. The resulting conformation, while being a purely computational prediction, hints at some properties which might contribute to the stability of the complex and support the functional interaction between TFEB and PAK6 that we have seen in our previous experiments. In particular, the C-terminal part of TFEB (in green) is placed in proximity of the same PAK6 site already linked to the functional interaction with sunitinib (model 4KS8). Moreover, S467 (highlighted in red) is located in the C-terminal part of TFEB, which we found to be crucial for the formation of a complex between the kinase and the transcription factor (Fig. 6F). Combining all these results together, we propose that one mechanism whereby PAK6 regulates TFEB nuclear translocation, occurs via formation of a heterotrimeric complex with phospho-14-3-3s (Fig. 6G).

### *Mbt* knockdown exacerbates DA neuron degeneration in *Drosophila* and PAK6 overexpression reduces phosphorylated α-syn in DA neurons of mutant LRRK2 mice

So far, we collected multiple lines of evidence that PAK6 activates autophagy through TFEB. Given the well-established role of impaired autophagy in neurodegeneration, including in PD, we next investigated whether PAK6 kinase activity could promote the autophagic clearance of the PD associated protein α-syn, which forms toxic neuronal aggregates in sporadic and genetic forms of PD [33]. To this end, we took advantage of *D. melanogaster* flies overexpressing the human α-syn in DA neurons [34] and crossed them with *mbt* KD flies. As figure 7 illustrates, we found that the downregulation of *mbt* strongly reduces the lifespan of α-syn-DA flies, indicating that the silencing of *mbt* may exacerbate the toxicity associated with α-syn overexpression (Fig. 7A). Supporting this hypothesis, *mbt* KD enhanced neuronal loss in flies overexpressing α-syn and increased the number of DA neurons positive for aggregated α-syn (Fig 7B).

**Figure 7.**
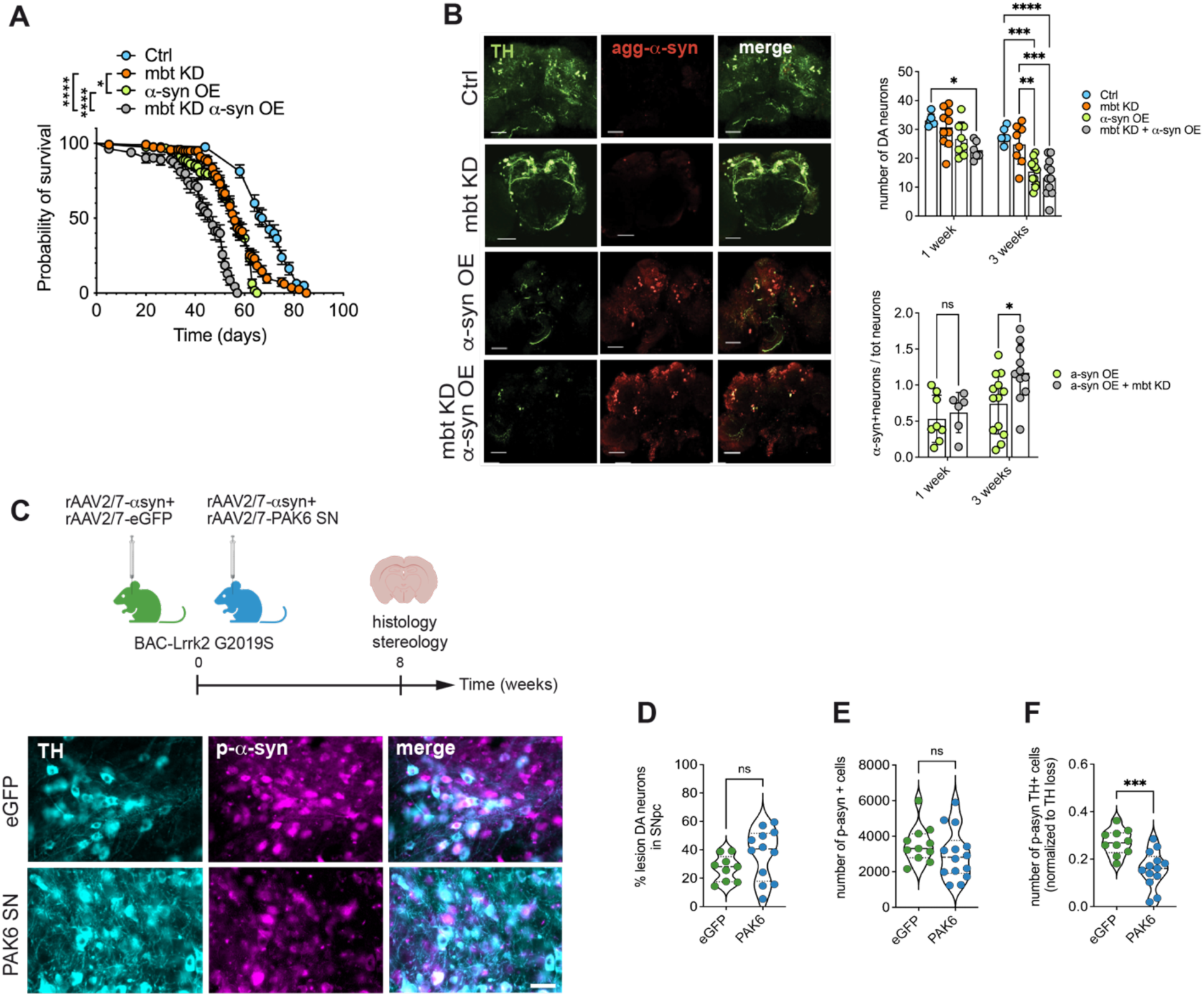
mbt/PAK6 affects alpha-synuclein pathology in dopaminergic neurons. (**A**) Lifespan of control, *mbt* KD, α-syn-OE, *mbt* KD + α-syn-OE. At least 60 flies per genotype were used in each experiment. Log-rank (Mantel-Cox) test (*mbt* KD vs *α-syn*-OE *p<0.05; *mbt* KD vs *mbt* KD + α-syn-OE ****p<0.0001; α-syn-OE vs *mbt* KD + α-syn-OE ****p<0.0001; control vs *mbt* KD****p<0.0001; control vs α-syn-OE****p<0.0001; control vs *mbt* KD + α-syn-OE****p<0.0001) (**B**) Immunohistochemistry of Drosophila brains stained with TH and aggregated α-syn (Scale bar 50 µm). Quantification of DA neuron number (top right) and α-syn-positive neurons (bottom right). Two-way ANOVA (DA neurons: interaction p=0.3085 F (3, 60) = 1.225; time p<0.0001 F (1, 60) = 35.63; genotype p<0.0001 F (3, 60) = 17.52; α-syn-positive neurons: interaction p=0.1386; F (1, 35) = 2.297; time: p=0.0109 F (1, 35) = 7.240; genotype: p=0.0524 F (1, 35) = 4.032) with Šídák’s multiple comparisons test (*p<0.05). (**C**) Experimental design (top), N= 11 (rAAV2/7 WT α-syn+eGFP) and N=14 (rAAV2/7 WT α-syn + PAK6 SN). TH and phospho-S129 α-syn immunohistochemical staining of the midbrain of BAC G2019S mice injected with rAAV2/7 WT α-syn combined with eGFP or PAK6 SN (bottom). Scalebar is 400µm. (**D**) Quantification of DA neurons in BAC G2019S mice injected with rAAV2/7 WT α-syn combined with eGFP or PAK6 SN (one sample t-test with outlier analysis, p>0.5). (**E**) Quantification of phospho-α-syn positive cells on the total midbrain cell population from BAC G2019S mice injected with rAAV2/7 WT α-syn combined with eGFP or PAK6 SN (one sample t-test with outlier analysis, p>0.5). (**F**) Quantification of phospho-α-syn positive cells in TH-positive neurons normalized to TH loss from BAC G2019S mice injected with rAAV2/7 WT α-syn combined with eGFP or PAK6 SN (one sample t-test with outlier analysis, ***p<0.01).

In parallel, we used the bacterial artificial chromosome (BAC)-*Lrrk2*-G2019S pre-symptomatic PD mouse overexpressing murine *Lrrk2* carrying the G2019S mutation [35], to assess the potential protective effect of PAK6 overexpression against dopaminergic neurodegeneration and α-syn pathology. Similar to a previous study on PAK4 [17] and based on the biochemical data reported in this study, we used the constitutively active form of PAK6 (S531N) to maximize the potential protective effect of the kinase. We first optimized the vector doses of 3xflag-PAK6^S531N^ and α-syn. We aimed for high expression levels of PAK6 to optimize potential beneficial effects and for α-syn levels leading to a mild neurodegenerative phenotype. A high level of PAK6 expression could be confirmed, as well as the lack of dopaminergic cell loss, based on tyrosine hydroxylase (TH) stereological analysis (Fig. S6A). To also optimize the α-syn vector dose and to assess whether expression of the pathogenic Lrrk2 G2019S variant affects α-syn-induced neurodegeneration, we injected rAAV2/7-α-syn in the *substantia nigra* (*SN*) of WT and Lrrk2 G2019S BAC mice. α-Syn expression was confirmed at 7 days and 8 weeks post-injection (p.i.) in all animals and dopaminergic neurodegeneration was assessed based on stereological analysis of TH-positive cells in the injected *versus* non-injected hemisphere (Fig. S6B). Although no significant difference was observed in dopaminergic cell death between α-syn-injected WT or BAC G2019S mice, the latter showed a trend towards increased sensitivity to α-syn-induced neurodegeneration (Fig. S6B). Having established the appropriate vector doses to obtain robust PAK6^S531N^ expression in the SN with no toxicity and mild α-syn-induced TH loss, we next investigated whether active PAK6 could offer protection against α-syn-induced neurodegeneration and/or pathology in a pathogenic Lrrk2 background. To this end, we injected BAC-Lrrk2 G2019S mice with human α-syn WT, combined with PAK6^S531N^ or eGFP as control (Fig. 7C). As shown in figure 7, ~30% DA neuronal loss could be observed in the eGFP control conditions, which represents the mild PD phenotype we aimed for (Fig. 7D). No significant difference in dopaminergic neurodegeneration could be observed between PAK6^S531N^-injected and eGFP injected mice (Fig. 7D). To evaluate the presence of α-syn pathology which could anticipate neuronal death, we determined the number of cells with α-syn phosphorylated at S129, which is considered the pathological form of the protein [36,37]. Interestingly, despite a similar α-syn pathological phosphorylation between PAK6^S531N^ and control conditions in the total cell population of the SN pars compacta (Fig. 7E), a closer look revealed a strong reduction in α-syn pathology in the TH-positive cell population in conditions of PAK6^S531N^ expression (Fig. 7C-F). This is a very interesting finding, revealing a modifying role of PAK6 selectively in TH neurons that are highly vulnerable in PD.

## Discussion

In this study, we identified a previously undisclosed role of the neuronal kinase PAK6 as an autophagy regulator via modulation of TFEB nuclear localization. Autophagy is a conserved cellular degradative mechanism ubiquitously performed in every cell type and precisely regulated to preserve a balanced homeostasis [38]. Autophagy correct function is particularly crucial for neuronal cells given their post-mitotic nature. Accordingly, dysregulation of neuronal autophagy is widely accepted as a contributor of the onset and progression of neurodegenerative diseases [1,8,9]. On this regard, induction of neuronal autophagy is considered a promising approach to clear dysfunctional organelles and protein aggregates in degenerating neurons [39]; however, its upregulation may have detrimental side-effects in other cells and tissues, where autophagic activation has been associated with different types of cancer [40]. For this reason, our work aimed at identifying a neuronal-specific modulation of the ALP. In fact, selective modulation of autophagy in neurons undergoing neurodegeneration should circumvent unwanted peripheral effects. Here we identified the brain enriched kinase PAK6 as a novel inducer of autophagy via TFEB. PAK6 belongs to the second group of the PAK protein family [22]. Noteworthy, PAK1 and PAK2 have already been demonstrated to contribute to the ALP regulation [11,15], whereas PAK4 depletion is associated with neuronal degeneration [12], supporting the link between PAK proteins family and the molecular pathways that control neuronal homeostasis and autophagy. Our interest in PAK6, which shares high degree of homology with the other members of the family, was mainly driven by its restricted pattern of expression. Indeed, in mammals the kinase is enriched in neurons, making it an ideal putative target to tune autophagy in this cell type [10].

Using a large variety of techniques and taking advantage of complementary models, we unravelled a novel function of PAK6 as an activator of TFEB, the master regulator of autophagy [2]. We obtained consistent results on PAK6-mediated TFEB nuclear translocation both *in vitro* using different immortalized human cell lines and *in vivo* in *C. elegans* suggesting that this cellular mechanism is highly conserved throughout the evolution. The degree of conservation frequently reflects the physiological relevance of a cell process and PAK6 may have evolved with its specific expression pattern as an important factor for the regulation of neuronal homeostasis by modulating the activity of TFEB.

We propose two possible parallel mechanisms of TFEB activation regulated by PAK6. A first one implies the PAK6-mediated phosphorylation of 14-3-3s, a family of seven proteins that acts as chaperones to modulate the localization and the activity of their binding partners [41]. This phosphorylation may promote 14-3-3s dissociation from TFEB, resulting in the nuclear translocation of the transcription factor. This scenario is supported by our previous study showing that 14-3-3s lose affinity for their substrates, including the PD-kinase LRRK2, when phosphorylated at Ser58/59 by PAK6 [19]. Such a mechanism falls within the class of canonical regulatory processes of TFEB, whose nuclear translocation is known to be prevented by the binding with the 14-3-3s [5,42]. A second mechanism entails a physical interaction between PAK6 and TFEB. Immunoprecipitation and fluorescence correlation imaging experiments showed that TFEB can indeed form a complex with constitutive active PAK6 provided that Ser467 of TFEB is phosphorylatable, a process that likely enhances and stabilizes the nuclear fraction of TFEB. Of interest, Ser467 belongs to a stretch of serine residues in the C-terminus of TFEB whose phosphorylation was shown to influence TFEB activity [30]. Specifically, Ser467 has been described to be the target of the energy sensor protein kinase AMPK, whose phosphorylation induces TFEB transcriptional activity [6]. Importantly, Ser467 is embedded in a PAKs consensus site, and our experimental data confirmed that phosphorylation of this aminoacid enhances the PAK6-mediated TFEB nuclear translocation. Future studies should investigate whether PAK6 directly phosphorylates TFEB at Ser467.

The correlation fluorescence microscopy data support a scenario where both mechanisms may concurrently occur within cells, implying that they would induce the nuclear translocation of TFEB with different kinetics. Furthermore, the biochemical data and the bioinformatic model suggest an additional layer of complexity where phosphorylated 14-3-3s take part of a tri-component protein complex with TFEB and PAK6, which results in increased fraction of the transcription factor in the nucleus. While additional studies are needed to precisely dissect the molecular aspects of these complex interactions, our findings pave the way for the existence of new mTORC1-independent mechanisms governing TFEB activation, with 14-3-3s acting as more complex players other than simple on-off switches.

As it appears for other studies, we were not able to identify a specific 14-3-3 isoform or subset of isoforms that preferentially bind TFEB upon PAK6 activation. One possibility is that the 14-3-3 isoform(s) binding TFEB is tissue/cell compartment/signal-specific, given the high degree of redundancy in the function and localization of these chaperones [41,43]. Furthermore, multiple isoforms may bind TFEB in different sites with specialized functions. Dissecting the identity of 14-3-3s isoforms involved in TFEB regulation would not only illuminate the intricate mechanisms governing TFEB’s activity but also pave the way for the development of innovative strategies to modulate its transcriptional function.

The findings that PAK6 activity affects the accumulation of pathological α-syn in *Drosophila* and in dopaminergic neurons of a mouse model of LRRK2-PD, provides strong evidence for considering PAK6 as an appealing therapeutic target to counteract neurodegeneration. Notably, we showed that PAK6 acts in the major pathway of TFEB regulation, i.e. downstream of mTORC1. Indeed, chemical inhibition of mTORC1 leads to an increase of PAK6 phosphorylation and activity. Considering that PAK6 knockout or knockdown does not block ALP but rather reduces its magnitude, we propose that the kinase acts as a fine tuner rather than an essential factor of ALP in neurons. This is also confirmed by the fact that *Pak6* KO murine models are viable [44,45]. Thus, increasing PAK6 activity in the brain, e.g through blood brain barrier penetrant compounds or via gene therapy, may be explored as a future therapeutic strategy for PD and other neurodegenerative disorders where autophagy activation is considered beneficial.

## Materials and methods

### Animals

#### Mouse lines

C57BL/6 LRRK2 wild-type and mouse Lrrk2 G2019S BAC (GS BAC) [B6.Cg-Tg(Lrrk2∗G2019S)2Yue/J] were provided by Jackson Laboratory. Housing and handling of mice were done in compliance with national guidelines. All animal procedures were approved by the Ethical Committee of KU Leuven (licence number LA1210579 to Veerle Baekelandt).

#### Drosophila melanogaster

*Drosophila* Uas-dsRNAi *mbt* (#29379), UAS-human-α-synuclein (#8146), yellow (#169), daughterless-Gal4 (#8641, da-Gal4), Repo-Gal4 (#90374), Elav-Gal4 (#458), and ple-Gal4 (#8848, TH-Gal4) lines were obtained from the Bloomington *Drosophila* Stock Center. All strains were reared on common cornmeal food in a humidified, temperature-controlled incubator at 25°C on a 12hr light/dark cycle.

#### Caenorhabditis elegans

The Bristol N2 (control animals), MAH240 [p*hlh-30*::*hlh-30*::*GFP* + *rol-6*(*su1006*)], and AML10 (otIs355 [p*rab-3*::*NLS*::*tagRFP*]; otIs45 [p*unc-119*::*GFP*]) strains were provided by the *Caenorhabditis* Genetics Center (CGC, University of Minnesota, Minneapolis, MN). The [p*rab-3*::*NLS*::*tagRFP*] strain was derived from AML10 by genetic crosses. Culture, maintenance, germline transformation, and genetic crosses were performed using standard techniques [46].

### Cell cultures, transfections and treatments

SH-SY5Y cells purchased from ATCC were cultured in a 1:1 mixture of Dulbecco’s modified. Eagle’s medium (DMEM, Life Technologies) and F12 medium, supplemented with 10% fetal bovine serum (FBS, Life Technologies). Cell lines were maintained at 37°C in a 5% CO_2_ controlled atmosphere. 0.25% trypsin (Life Technologies), supplemented with 0.53mM EDTA, was employed to generate subcultures. Stable cell lines overexpressing PAK6 wild type were generated as described in [18]. Briefly, the cDNA sequence encoding PAK6 was cloned into the lentiviral plasmid pCHMWS-MCS-ires-hygro. 500 ug/ml hygromycin was utilized for selection, while 100 ug/ml for maintenance.

Human Embryonic Kidney 293T (HEK293T) and HeLa cells were cultured in DMEM supplemented with 10% FBS (Life Technologies). 1 % trypsin was employed to detach cells and split them. HEK293T and HeLa cells were transfected with plasmid DNA using polyethylenimine (PEI, Polysciences) for 48 hours according to the manufacturer’s instructions.

### Cell lysis and western blot

Fly tissues or cells were lysed in 50 mM Tris-HCl pH 7.5, 1 mM sodium orthovanadate, 50 mM sodium fluoride, 10 mM β-glycerophosphate, 5 mM sodium pyrophosphate, 1% Triton X-100, 1 mM EGTA, and 270 mM sucrose buffer added with protease inhibitors (Roche) and incubated on ice for 30 minutes. Subsequently, lysates were cleared by centrifugation at 20000 x g for 30 minutes at 4°C. Supernatants were used. Total proteins amount was quantified by using Pierce™ BCA Protein Assay Kit (Thermo Fisher Scientific). For each sample, 30-50 μg of protein were loaded on or ExpressPlus™ PAGE 4–20% gels (GenScript), in MOPS running buffer or 7.5%, 10% Tris-glycine polyacrylamide gels in SDS/Tris-glycine running buffer, according to the size resolution required. Precision Plus molecular weight markers (Bio-Rad) were used for size estimation. The resolved proteins were transferred to polyvinylidenedifluoride (PVDF) membranes (Bio-Rad) or nitrocellulose membranes (Whatman Products), through a Trans-Blot® Turbo™ Transfer System (Bio-Rad). Membranes were subsequently blocked in Tris-buffered saline plus 0.1% Tween (TBS-T) plus 5% non-fat milk for 1 hour at room temperature (RT) and then incubated over-night at 4 °C with primary antibodies diluted in TBS-T plus 5% non-fat milk. Membranes were then washed in TBS-T (3×10 minutes) at RT and subsequently incubated for 1hour at RT with horseradish peroxidase (HRP)-conjugated secondary antibodies. Blots were then washed in TBS-T (3×10 min) at RT and rinsed in TBS-T, and immunoreactive proteins were visualized using Immobilon® Forte Western HRP Substrate (Merck Millipore) at the Imager Chemi Premium (VWR International). Densitometric analysis was carried out using the Image J software. Antibodies utilized for western blot: fly samples: mouse β-actin (Sigma Prestige); rabbit LC3 (Thermo Fisher Scientific, PA1-16930); goat Anti PAK6 (Novus Biological, AF4265); mouse Hsp-70 (Stressgen Enzo Life science, SPA810). Human cell line samples: mouse anti-β-Actin (Sigma-Aldrich, A1978); rabbit anti-PARP (Cell Signalling Technology, 9542S); anti-Flag M2-Peroxidase (Sigma-Aldrich, A8592); anti GFP (Roche-Sigma, 11814150001); rabbit Anti-Phospho-PAK4/5/6 (pSer474) (Sigma-Aldrich, SAB4503964); rabbit Anti-Phospho-14-3-3 (pSer58) (Abcam, ab30554); Rabbit anti-TFEB (Bethyl Laboratories, A303-673A); mouse anti-LAMP1 (H4A3) (Santa Cruz Biotechnology, sc-20011); rabbit anti-p62 (Abcam, ab109012)

### Co-immunoprecipitation assay

Cells were harvested at 48 h post transfection and lysed in buffer containing 20 mM Tris-HCl pH 7.5, 150mM NaCl, 1mM EDTA, 2.5mM sodium pyrophosphate, 1mM beta-glycerophosphate, 1mM sodium orthovanadate, 1% v/v Tween®, 20 and 1% of protease inhibitor cocktail (Roche). The resuspended cells were kept on ice for 30 minutes and then centrifuged for 30 minutes at maximum speed at 4°C. The supernatants were then incubated overnight with GFP-Trap® (Chromotek). The GFP-tagged protein bound to resins were then washed 10 times with the following Washing Buffers (WB), two washes for each WB: WB1 (20mM Tris-HCl, pH 7.5, 500mM NaCl, 1% v/v Tween® 20); WB2 (20mM Tris-HCl, pH 7.5, 300mM NaCl, 0,5% v/v Tween® 20); WB3 (20mM Tris-HCl, pH 7.5, 150mM NaCl, 0,5% v/v Tween® 20); WB4 (20mM Tris-HCl, pH 7.5, 150mM NaCl, 0.1% v/v Tween® 20); WB5 (20mM Tris-HCl, pH 7.5, 150mM NaCl, 0.02% Tween® 20). Immunoprecipitates were resuspended in sample buffer. Between 15 and 30 μg of proteins were resolved on 4-20% gels.

### Cell fractionation

Cell fractionation was performed with the NE-PER™ Nuclear and Cytoplasmic Extraction Reagents (Thermo-Scientific) kit following the manufacturer’s instructions. Cytoplasmatic and nuclear samples were resuspended in sample buffer and loaded in 10% Tris-glycine polyacrylamide gels as previously described for western blots.

### Immunocytochemistry and confocal imaging

HeLa or SH-SY5Y cells were cultured onto 12mm glass coverslips (Thermo-Scientific) coated with Poly-L-Lysine (Sigma-Aldrich). Cells were washed in PBS and fixed with 4% w/v Paraformaldehyde (PFA) for 20 minutes at RT. After cell permeabilization in PBS plus triton 0,1% for 20 minutes at RT and a blocking step performed in 3% BSA diluted in PBS for 30 minutes at RT, cells were stained with the appropriate primary antibody diluted in PBS plus 3% BSA for 1 hour at RT. Subsequently, cells were washed three times in PBS and incubated with the secondary antibody Alexa Fluor® 488 conjugated (Life Technologies) or Alexa Fluor® 568 conjugated (Life Technologies). Before mounting the coverslips on glass slides, cells were incubated with Hoechst 33258 (Invitrogen) diluted in PBS for 5 minutes. Images were acquired with a LeicaSP5 confocal microscope (Leica Microsystems) and quantified using ImageJ. The following antibodies were used: mouse anti-Flag F7425 (Sigma-Aldrich, F7425); rabbit anti-Phospho-PAK4/5/6 (pSer474) (Sigma-Aldrich, SAB4503964).

### TFEB nuclear translocation

The analysis of the nuclear translocation of endogenous TFEB has been performed as follow: 10k cells per well were plated in 96 well plates (PerkinElmer, 6055302) and transfected in suspension using Lipofectamine LTX (Invitrogen, 15338) according to manufacturer instructions. Twenty-four hours after transfection, the plate was acquired with Opera Fenix (PerkinElmer). Analysis was performed with Harmony software (PerkinElmer). Transfection-positive cells were first selected and then cytoplasmic and nucleus intensities were calculated. TFEB translocation is expressed as the ratio of nuclear intensity to cytoplasmic intensity.

The nuclear translocation of overexpressed TFEB was done in cells transfected with plasmid DNA of TFEB-eGFP and Flag-PAK6 mutants for 48 hours. To induce TFEB nuclear translocation and autophagic activity cells were treated with 2.5 μM of the specific inhibitor of mTORC1 Torin1 (MedChemExpress, HY-13003) for 90 minutes. Cells were then fixed with 4% PFA, stained for flag and Dapi and analyzed via confocal microscopy. TFEB translocation is expressed as the ratio between TFEB intensity in the nucleus minus TFEB intensity in the cytoplasm to total TFEB intensity, and correlated to PAK6 intensity.

### GCase enzymatic assay

GCase activity was measured using a protocol similar to the one described in [47]. Briefly, fly lysates were obtained, and the total amount of proteins was quantified as previously described. 20 μl of lysate were diluted in 40 μl of a solution of citrate phosphate buffer (0.1 M citric acid, 0.2 M solution dibasic sodium phosphate), 0,2% sodium taurodeoxycholate hydrate and 3 mM 4-methylumbelliferyl β-D-glucopyranosidase (Sigma-Aldrich, M3633) and incubated at 37°C for 90 minutes. The reaction was stopped by adding ice-cold glycine buffer (0.2 M solution Glycine, 0.2 M NaOH) and fluorescence was detected on a multilabel plate reader Victor (Perkin Elmer) at excitation 360 nm and emission 440 nm. Final GBA activity values were calculated normalizing the value obtained to the total amount of protein present in the sample.

### Scanning fluorescence correlation spectroscopy (FCS) microscopy experiments

All dual-channel line scan FCS measurements were performed on an Olympus FV3000 confocal laser scanning microscope (CLSM) with a 60x water-immersion objective (1.2 NA) and temperature as well as CO2 control (37° in 5% CO2). The eGFP and mCherry plasmids were excited by solid-state laser diodes operating at 488 nm and 561 nm, respectively, and the resulting signal was directed through a 405/488/561 dichroic mirror to two internal GaAsP photomultiplier detectors set to collect 500–540 nm and 600–700 nm, respectively. The dual-channel line scan FCS measurements (n = 100,000 lines) were acquired with: (1) a 64-pixel line size at a digital zoom that resulted in a line length of 5.3 μm and a pixel size 83 nm, and (2) a 8 μs pixel dwell time that resulted in a line time of 1.624 ms (our sampling frequency).

### Drosophila melanogaster assays

#### Life span

Adult males (1-3 day-old) were collected under brief CO_2_ exposure and transferred into new tubes containing standard food (20 flies/vial). Flies were transferred to fresh food vials every 3-4 days and the number of dead flies was counted daily. The percentage of survival was calculated at the end of the experiments.

#### Lysotracker staining

Adult males (1-3 day-old) brains were dissected in phosphate-buffered saline (PBS) followed by 15 minutes of incubation with 200 nM LysoTracker™ Red DND-99 Red probe (Thermofisher Scientific, L7528) in the dark. Subsequently, brains were washed trice in PBS for 5 minutes, mounted in Mowiol ® 4-88 (Calbiochem, 475904), and immediately analyzed. Z-stack images were acquired on a LeicaSP5 confocal microscope (Leica Microsystems). The area of the brain occupied by LysoTracker staining and the number of acidic compartments were quantified by ImageJ.

#### *Drosophila* dopaminergic neurons and alpha-synuclein analysis

Brains from 1-3 days old male flies were labelled with anti-TH-and anti-aggregated α-syn. In brief, brains were dissected and fixed in 0.4% paraformaldehyde (PFA) for 1h at RT. After permeabilization in 1% triton dissolved in PBS for 10 minutes and blocking in 1% BSA, 0,3% triton dissolved in PBS for 1 hour at RT, brains were stained overnight with primary antibodies diluted in PBS plus 0,3% triton, and 0,1% BSA at 4°C. Brains were then incubated with the fluorophore-conjugated secondary antibodies for 2 hours at RT. Finally, brains were mounted in Mowiol® 4-88 (Calbiochem, 475904), and analyzed. Images were acquired using a LeicaSP5 confocal microscope (Leica Microsystems), the number of posterior dopaminergic neurons and α-syn-containing DA neurons were counted manually in each brain hemisphere. The following antibodies were used for this experiment: rabbit anti-tyrosine hydroxylase (Merkmillipore, AB152); mouse anti-aggregated-α-synuclein, clone 5G4 (Sigma Aldrich, MABN389).

### *C. elegans* HLH-30 nuclear translocation assay

The cDNA corresponding to the wild-type or S531N human *PAK6* alleles were subcloned into the pBy103 vector (a kind gift from E. Di Schiavi, IBBR-CNR, Naples) to be under the control of the *unc-119* promoter, which drives pan-neuronal expression, and were injected at 30 ng/µl to generate multi-copy extrachromosomal arrays. The pJM371 plasmid (p*elt-2*::*NLS*::*tagRFP*) (a kind gift from E. Di Schiavi, IBBR-CNR, Naples), which drives RFP expression in intestinal cell nuclei, was used as co-injection marker, and injected at 30 ng/µl. Isogenic lines that had lost the transgene were cloned separately and used as controls. The subcellular localization of HLH-30 prior to and following short-term (120 and 180 minutes) or long-term (overnight) starvation was investigated in p*hlh-30*::*hlh-30*::*GFP*;p*rab-3*::*NLS*::*tagRFP* animals expressing or not the wild-type or mutant allele of hPAK6. Fluorescent microscopy analysis was performed using an Eclipse Ti2-E microscope (Nikon Europe, Florence, Italy) equipped with DIC optics on live animals mounted on 2% agarose pads containing 10 mM sodium azide as anesthetic. Starvation was obtained by moving young adult hermaphrodites to NGM plates without food for two or three hours, or overnight.

### *In vivo* AAV-injections and histology in mouse

Two µl of rAAV CamKII 0.4-intron-3flag-PAK6 S531N at 5.6 E11 GC (genome copies)/ml was injected in the substantia nigra (SN) of WT mice to confirm transgene expression and the lack of toxicity. Mice were sacrificed 8 weeks post injection (p.i.) and stained for PAK6 and TH. To optimize the α-syn vector dose and to assess whether expression of the pathogenic Lrrk2 G2019S variant affects α-syn-induced neurodegeneration, 2 µl rAAV2/7 CMVie synapsine1 α-syn at a concentration of 2.00 E12 GC/ml were injected in the SN of WT and Lrrk2 G2019S BAC mice. To take into account the higher vector load in the co-injection experiment of α-syn with PAK6 S531N, 2µl of CamKII 0.4 intron 3xflag-eGFP (1.4 E12 GC/ml) were injected together with the α-syn vector to obtain comparable experimental conditions. Nine mice were sacrificed 7 days p.i. (i.e. 4 BAC Lrrk2 G2019S mice and 5 littermates) to confirm α-syn expression and the lack of vector toxicity. Nineteen mice were sacrificed 8 weeks p.i. (i.e. 9 BAC Lrrk2 G2019S and 10 WT mice). To evaluate PAK6 S531N protective activity in the LRRK2 mouse model injected with α-syn, Lrrk2-G2019S BAC transgenic mice were injected with human α-syn WT (1.44 E12 GC/ml), combined with PAK6 S531N (5.6 E11 GC/ml) or eGFP (6.9 E11 GC/ml) as control. Mice were sacrificed 8 weeks p.i. (i.e. 12 eGFP- and 13 PAK6 SN-injected mice).

### Computational analysis

#### Co-expression analysis

We downloaded from the Human Protein Atlas the “RNA consensus tissue gene data”, including transcriptomics measurements from HPA (https://www.proteinatlas.org/about/assays+annotation#hpa_rna) and GTEx (https://www.proteinatlas.org/about/assays+annotation#gtex_rna). After loading the matrix in the R statistical environment, we extracted the normalized nTPM values and averaged repeated gene measures for the same tissue. We computed the Pearson correlation among all expression profiles, and for each of PAK4, PAK5 and PAK6 we identified the 50 most co-expressed genes. We finally plotted an heatmap for each gene group, representing the expression levels in the logarithmic scale.

#### *In silico* docking model

We have employed a computational approach to model a putative tripartite complex among phospho-TFEB, phospho-14-3-3 and PAK6. Our specific goal was to predict how their interactions might lead to the formation of a stable complex. As a first step, we downloaded a PAK6 model (accession Q9NQU5) from the Swiss-model database, while a crystal structure for 14-3-3 in complex with TFEB was obtained from PDB (accession 6A5Q). The latter model only includes the binding motif for TFEB, so we applied the AlphaFold 2 method (using the more sensitive “full_dbs” preset) to obtain an *in silico* prediction for its entire structure. We then submitted these results to the ClusPro webserver to evaluate the docking among the different proteins. As this software requires a pair of structures as an input, we first computed the docking among TFEB and 14-3-3, and we later considered PAK6. For each run, ClusPro provides ten solutions with the lowest energy among all the conformations it has computed. Every conformation is in principle associated with a numeric “Model Score”, but the authors of the method advise against using such scores for ranking the results. We therefore looked for a solution in which S58/59 on 14-3-3s was sufficiently close to PAK6 to make its phosphorylation possible.

## Acknowledgments

This work was supported by the Michael J Fox Foundation for Parkinson Research (EG and VB, grant number 15593), the University of Padova BIRD funding (EG).

Bioinformatics analyses were performed using the high-performance computing system funded by the University of Padova Strategic Research Infrastructure Grant 2017: “CAPRI: Calcolo ad Alte Prestazioni per la Ricerca e l’Innovazione”.

## ABBREVIATIONS

AD: Alzheimer Disease
ALP: autophagic lysosomal pathway
α-syn: alpha-synuclein
AMPK: AMP-activated protein kinase
CLEAR: Coordinated Lysosomal Expression and Regulation
DA: dopamine
FCS: fluorescence correlation spectroscopy
GCase: Glucocerebrosidase
KD: knockdown
KO: knockout
LRRK2: leucine-rich repeat kinase 2
mbt: mushroom bodies tiny
mTORC1: mechanistic target of rapamycin complex 1
OE: overexpressing
PAK4: p21 activated kinase 4
PAK5: p21 activated kinase 5
PAK6: p21 activated kinase 6
PD: Parkinson Disease
WT: wild type

## Notes

### Competing Interest Statement

The authors have declared no competing interest.

### Summary of Updates

Figure 2 has been replaced to add a revised version of the data.

