## Supplementary Materials for "PAK6 promotes neuronal autophagy by regulating TFEB nuclear translocation"

**Figure S1. Phenotypic characterization of ubiquitous and cell-specific *mbt* downregulation.**

Lifespan of flies as compared to control individuals (Ctrl). Analysis has been performed by using (**A**) total body (da) *mbt* KO flies, (**B**) glia (repo) *mbt* KO flies and (**C**) neuronal (elav) *mbt* KO flies compared with their control genotype. At least 40 flies per genotype were used in each experiment. Log-rank (Mantel-Cox) test, ns (non-significant), ****$p<0.0001$. (**D**) Climbing ability of total body control versus *mbt* KD flies. Five independent climbing evaluations were performed and the data were analyzed using unpaired t-test (**** $p < 0.0001$). (**E**) Ratio of fly eclosion. Six independent analysis were performed and analyzed using t-test (****$p<0.0001$). (**F**) Western blot analysis to confirm mbt downregulation in the whole-body. Data were analyzed using unpaired t-test from n=3 independent pools of flies (** $p < 0.01$). (**G**) Western blot of Atg8a levels in total body *mbt* knockdown. Data were analyzed using a t-test from n=3 independent pools of flies (* $p < 0.05$). (**H**) GCase enzymatic assay of fly heads downregulating mbt in total body and matching controls. N=5-6 biological replicates. Statistical significance was determined by unpaired t-test (* $p < 0.05$).

**Figure S2. Co-expression analysis**

Top-50 co-expressed genes from Human Protein Atlas the “RNA consensus tissue gene data” for PAK4 (**A**), PAK5 (**B**) and PAK6 (**C**).

**Figure S3. Constitutively active PAK6 induces nuclear translocation of GFP-TFEB *in vitro*.**

(**A**) TFEB nuclear translocation assessed in Hela cells cotransfected with GFP-TFEB and Flag-PAK6 WT, K436M (kinase dead) or S531N (constitutively active) (Scale bar 10 μm). Cells transfected with TFEB WT only were treated with 2.5 μM of Torin1 for 90 minutes as a positive control of TFEB nuclear translocation. Between X and Y cells per genotype were analyzed.

(**B**) Quantification of (**A**). Number of cells analyzed: Gus=52; PAK6-WT=85; PAK6-K436M=23; PAK6-S531N=81; Torin-1=107. Statistical significance was determined using one-way ANOVA with Tukey's multiple comparisons test (****$p<0.0001$).

(**C**) Scattered plots of TFEB nuclear translocation index ($TFEB_{cyto}$ - $TFEB_{nucleus}$ / $TFEB_{cyto}$ + $TFEB_{nucleus}$) against fluorescence intensity of PAK6 (or GUS control) to correlate the entity of translocation with the degree of overexpression.

**Figure S4. C. elegans and HLH-30 and human TFEB alignment and colocalization**

(**A**) Amino acid sequence alignment of *C. elegans* HLH-30 and human TFEB. * = identity between amino acids, : = conserved substitution, . = semi-conserved substitution.

(**B**) While HLH-30/TFEB localizes in the cytoplasm of neurons of the ventral nerve cord in well-fed p*hlh-30*::*hlh-30*::*GFP*;p*rab-3*::*NLS*::*tagRFP* animals (see Fig. 4C), it migrates into the nucleus in the absence of food (overnight starvation). Scale bar, 10 μm. Ninety neurons belonging to ten animals were tested.

**Figure S5. Schematic representation of the FCS applied to the live cell imaging.**

Image examples for CH1 and CH2 (EGFP and mCH signals) as they are acquired to perform FCS image analysis, followed by the two boxes representing the two type of correlation analysis performed, i.e. ACF and pCF. ACF analysis of the fluorescence signal measured in the cell nucleus was performed for the CH1:EGFP signal (green curve) and for the CH2:mCH signal (red curve). Cross correlation function (CC) was also evaluated (yellow curve). pCF analysis of the CH1 and CH2 channels from the cysotol to the nucleus (blu curves) and from the nucleus to the cytosplasm (orange curves); moreover, cross pCF was calculated for the complex formed by TFEB and PAK6 (CC) in both directions.

**Figure S6. Diffusion coefficient values for TFEB, PAK6 and the TFEB-PAK6 complex.**

ACF analysis allow the estimation of diffusion coefficients D (here expressed as logarithm, in Log $\mu m^2/s$) for the different proteins (wild type and mutants) or protein complexes. D values were calculated within the nucleus and in the cytoplasm, showing a two-component behaviour for TFEB (D1 and D2), and a single component behaviour for both PAK6 and the TFEB-PAK6 complex. ACF curves did not allow to calculated the D values for CC in TFEB WT PAK6 SN, TFEB SA PAK6 WT and TFEB SA PAK6 SN were expressed.

**Figure S7. Diffusion coefficients as evaluated across cystoplasm/nuclear barrier using pCF**

Pair correlation functions calculated from multiple line scan acquisitions that record the fraction of molecules that are transported from the cytoplasm to nucleus (C-N) versus nucleus to cytoplasm

(N-C) for the cells expressing the different combination of WT and mutant TFEB and PAK6 proteins. Curves and the associated diffusion coefficients are reported.

**Figure S8.**

**(A)** TH or Flag immunohistochemical staining of the midbrain of WT mice 8 weeks after injection in the right hemisphere of rAAV2/7 encoding human WT α-syn or 3flag-PAK6 SN. On the right, Stereological quantification of TH-positive cells in the SN pars compacta of mice expressing human WT α-syn or 3flag-PAK6 SN. Scalebar 1mm.

**(B)** α-syn immunohistochemical staining of the midbrain of WT or BAC G2019S mice 7 days after injection of rAAV2/7 encoding human WT α-syn. TH immunohistochemical staining of the midbrain of WT or BAC G2019S mice 7 days or 8 weeks after injection. Right: relative dopaminergic cell loss based on stereological quantification of TH-positive cells in the SN pars compacta of mice expressing human WT α-syn. Scalebar 400µm.

**Figure S1**

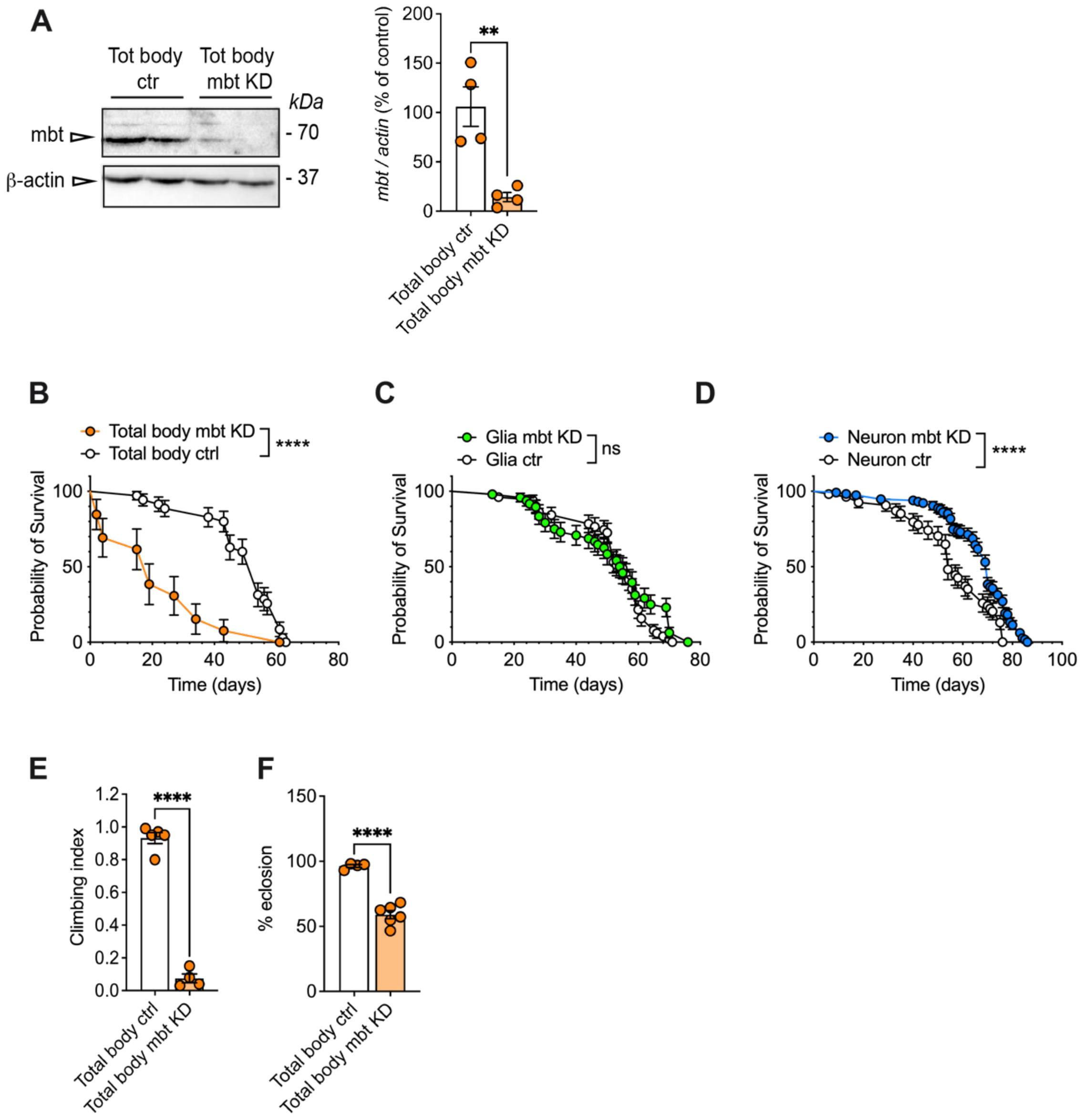
A
Tot body ctr
Tot body mbt KD
kDa
mbt
β-actin
- 70
- 37
mbt / actin (% of control)
**
Total body ctr
Total body mbt KD
B
Total body mbt KD
Total body ctrl
****
Probability of Survival
Time (days)
C
Glia mbt KD
Glia ctr
ns
D
Neuron mbt KD
Neuron ctr
****
E
Climbing index
****
Total body ctrl
Total body mbt KD
F
% eclosion
****

### Figure S2

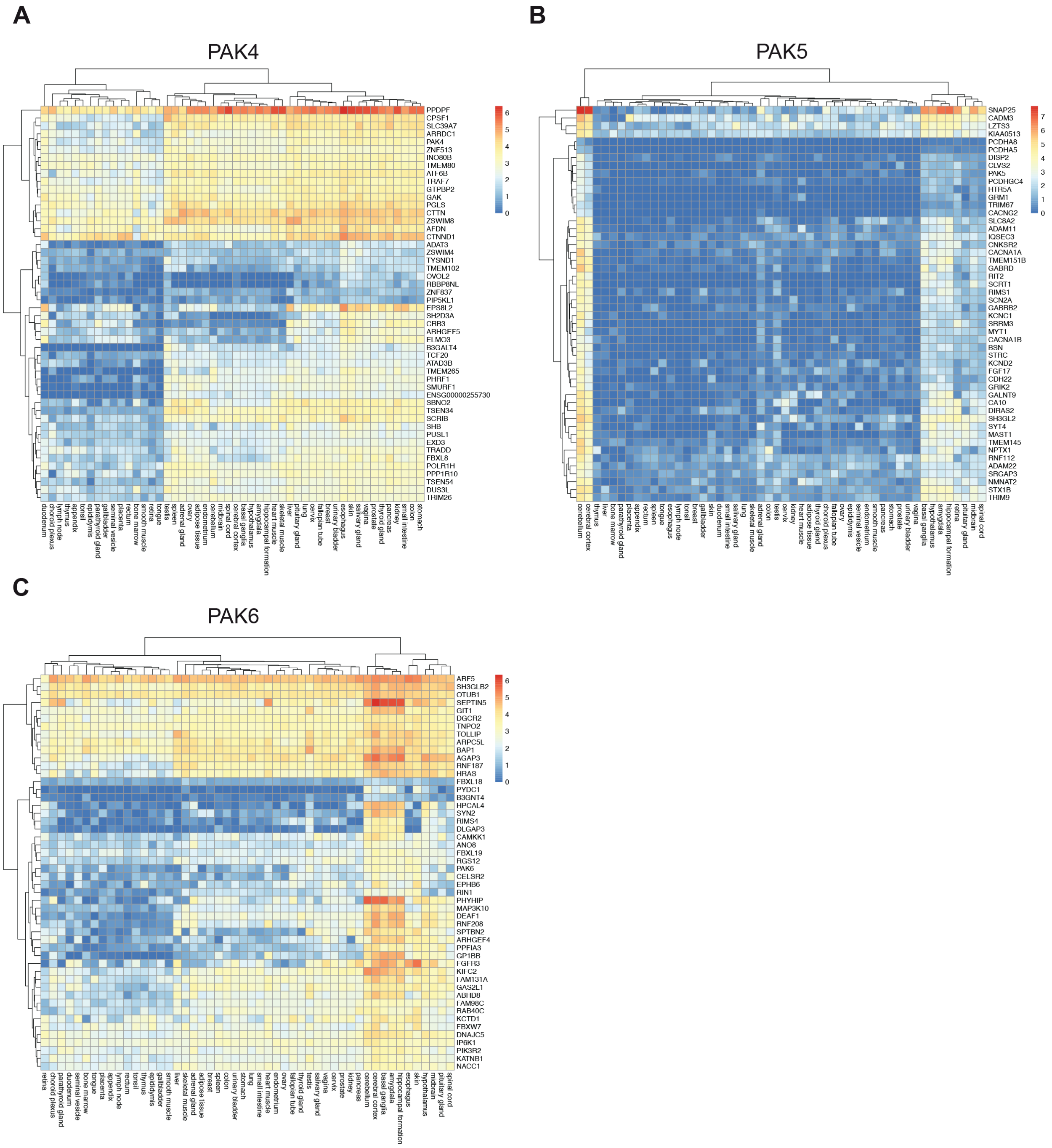
A
PAK4
PPDPF
CPSF1
SLC39A7
ARRDC1
PAK4
ZNF513
INO80B
TMEM80
ATF6B
TRAF7
GTPBP2
GAK
PGLS
CTTN
ZSWIM8
AFDN
CTNND1
ADAT3
ZSWIM4
TYSND1
TMEM102
OVOL2
RBBP8NL
ZNF837
PIP5KL1
EPS8L2
SH2D3A
CRB3
ARHGEF5
ELMO3
B3GALT4
TCF20
ATAD3B
TMEM265
PHRF1
SMURF1
ENSG00000255730
SBNO2
TSEN34
SCRIB
SHB
PUSL1
EXD3
TRADD
FBXL8
POLR1H
PPP1R10
TSEN54
DUS3L
TRIM26
B
PAK5
SNAP25
CADM3
LZTS3
KIAA0513
PCDHA8
PCDHA5
DISP2
CLVS2
PAK5
PCDHGC4
HTR5A
GRM1
TRIM67
CACNG2
SLC8A2
ADAM11
IQSEC3
CNKSR2
CACNA1A
TMEM151B
GABRD
RIT2
SCRT1
RIMS1
SCN2A
GABRB2
KCNC1
SRRM3
MYT1
CACNA1B
BSN
STRC
KCND2
FGF17
CDH22
GRIK2
GALNT9
CA10
DIRAS2
SH3GL2
SYT4
MAST1
TMEM145
NPTX1
RNF112
ADAM22
SRGAP3
NMNAT2
STX1B
TRIM9
C
PAK6
ARF5
SH3GLB2
OTUB1
SEPTIN5
GIT1
DGCR2
TNPO2
TOLLIP
ARPC5L
BAP1
AGAP3
RNF187
HRAS
FBXL18
PYDC1
B3GNT4
HPCAL4
SYN2
RIMS4
DLGAP3
CAMKK1
ANO8
FBXL19
RGS12
PAK6
CELSR2
EPHB6
RIN1
PHYHIP
MAP3K10
DEAF1
RNF208
SPTBN2
ARHGEF4
PPFIA3
GP1BB
FGFR3
KIFC2
FAM131A
GAS2L1
ABHD8
FAM98C
RAB40C
KCTD1
FBXW7
DNAJC5
IP6K1
PIK3R2
KATNB1
NACC1

**Figure S3**

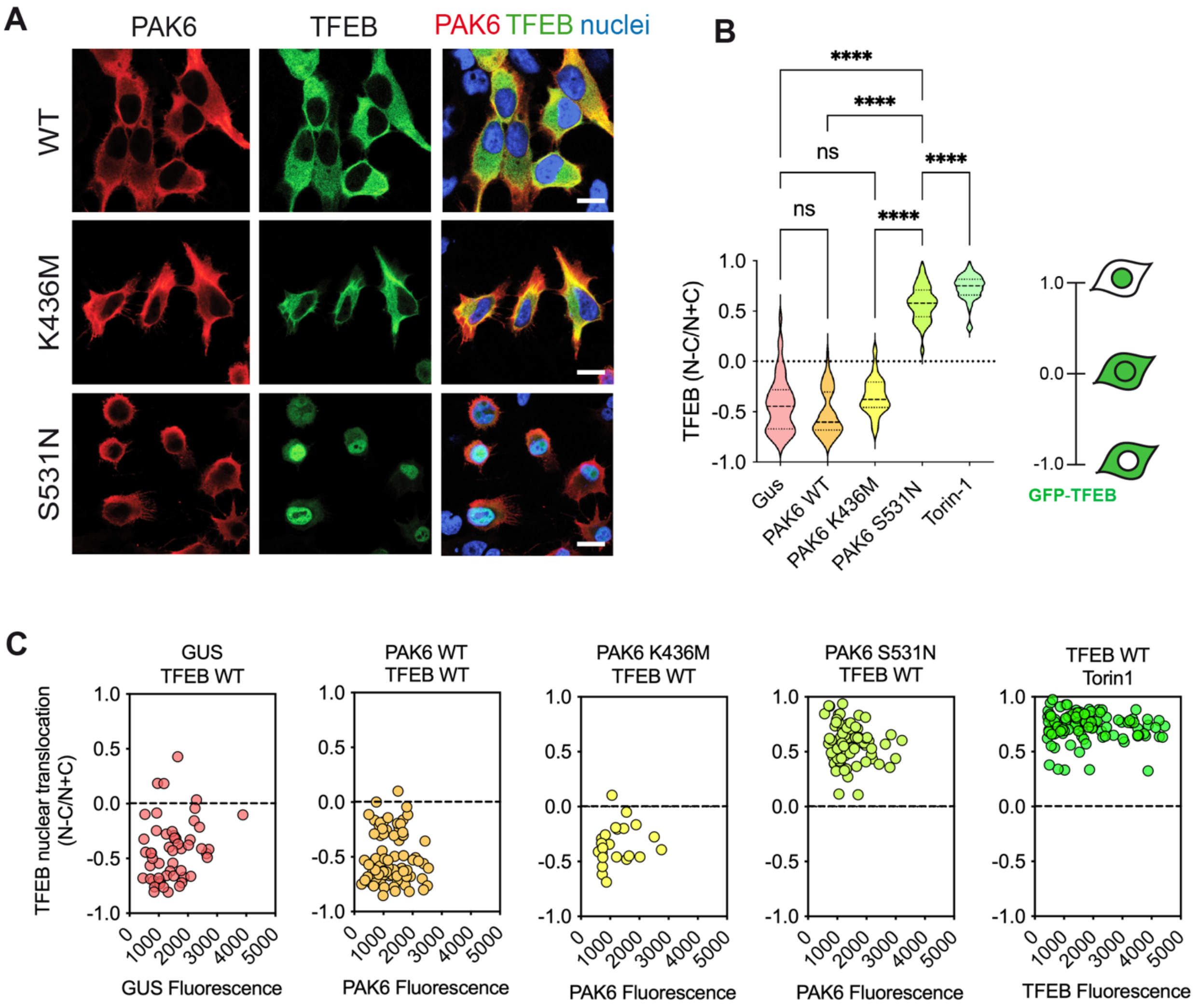
A
PAK6
TFEB
PAK6 TFEB nuclei
WT
K436M
S531N
B
****
****
ns
****
ns
****
TFEB (N-C/N+C)
1.0
0.5
0.0
-0.5
-1.0
Gus
PAK6 WT
PAK6 K436M
PAK6 S531N
Torin-1
GFP-TFEB
C
GUS
TFEB WT
PAK6 WT
TFEB WT
PAK6 K436M
TFEB WT
PAK6 S531N
TFEB WT
TFEB WT
Torin1
TFEB nuclear translocation
(N-C/N+C)
GUS Fluorescence
PAK6 Fluorescence
PAK6 Fluorescence
PAK6 Fluorescence
TFEB Fluorescence
0
1000
2000
3000
4000
5000

**Figure S4**

**A**

```
TFEB            -------MASRIGLRMQLMREQAQQEEQRERMQQQAVMHYMQQQQQQQQQLGGPPTPAI   53
HLH-30          MIRQLNSPGGGGGLGLNNPRAQQPPGAQQQ-----------QQPQQAQQQFYDDEPYQAN   49
                        ..  ** ::  * *     *::           ** ** ***  .. *  *

TFEB            NTPVHFQSPPP-----VPGE-----------------VLKVQSYLENPTSYHLQQSQHQK   91
HLH-30          ASQFRFGAGKSMEQRRETGNLIPIAQRSMGSTSTPFGSAPTQSYFGGGSSGAAL-SSPRK   108
                 : .:* :          *:                    .***: . :*     *. :*

TFEB            VREYLSETYGNKFAAHISPAQGSPKPPPAASPGVRAGHVLSSSAGNSAPNSPMAMLHIGS   151
HLH-30          MQQTHQMLF-----GNIQPPRGSPPSD-----GSDKIHRFGES---PTPG------GVGG   149
                :::  .  :     .:*.* :***        *    * :..*    :*.       :*.

TFEB            NPERELDDV-IDNIMRLDDVLGY---------IN--PEMQMPNTLPLSSSHLNVYSSD-P   198
HLH-30          VFGTELDDLIIDELMGMEDDQRMRPGATRPMTIGGEKTMSMARPIPGASSRAGSGHSGSP   209
                    ****: **::* ::*             *.    *.* . :* :**: .   *. *

TFEB            -----QVTASLVGVTSSSCPADLTQ------------KRELTDAESRALAKERQKKDNHN   241
HLH-30          ITIPNAMSNNFRQVVSSSAPTSSIDIEKMIGAVSNGGGNSGGDNDPEDYYRDRRKKDIHN   269
                      :: .:  *.***.*:.  :             ..  * : .   ::*:*** **

TFEB            LIERRRRFNINDRIKELGMLIPKANDLDVRWNKGTILKASVDYIRRMQKDLQKSRELENH   301
HLH-30          MIERRRRYNINDRIKELGQMLPKNTSEDMKLNKGTILKASCDYIRVLQKDREQAMKTQQQ   329
                :******:********** ::** .. *:: ********* **** :*** ::: : :::

TFEB            SRRLEMTNKQLWLRIQELEMQARVHGLPTTSPSGMNMAELAQQVVKQEL---PSEEGPGE   358
HLH-30          QKSLESTAHKYADRVKELEEMLARQGVQVPPSHLPPIPKVIERPIKQEIDESPPNHTPTG   389
                .: ** * ::   *::***     :*: .       : :: :: :***:   * :. *

TFEB            ALMLGA----EVPD-PEPLPALPPQAPLPLPTQPPSPFHHLDFSHSLSFGGREDEGPPGY   413
HLH-30          SFVSSSGFLSEVTNNTAAMQITSP-----NDSR-PNNFMNNS-APSDSFFSVGSASPPDY   442
                ::: .:    ** :    :    *       :: *. * : . : * ** .  . .**.*

TFEB            PEPLAP-----GHGSPFPSLSKKDLDLMLLDDSLLPLASDPLLSTMSPEASKAS--S---   463
HLH-30          RTSSGTASWKLPGSNAFSDLMMDDLNPMM--------NGDPLISSAGAHPSPHFHSSQMS   494
                    .        .. * .*  .**: *:         .***:*: . . *     *

TFEB            -----RRSSFSMEE--------GDVL----     476
HLH-30          PDIHWDASGFSPDPINTQQSNSGHYHMDFS     524
                       *.** :         *.
```

Yellow: basic helix loop helix motif
Grey: Ser122, Ser134, Ser138, Ser142
(important residues in the regulation of TFEB)
Red: Activation domain
Magenta: Ser211 (most conserved serine among those phosphorylated by mTORC1 and site of interaction between TFEB and 14-3-3s)
Cyan: Nuclear localization signal
Green: Zip domain
Blue: C-terminal serines

**B**

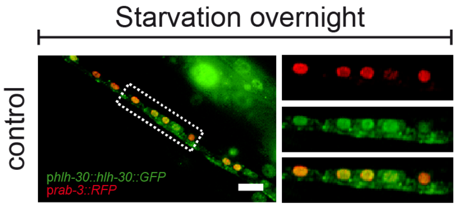

**Figure S5**

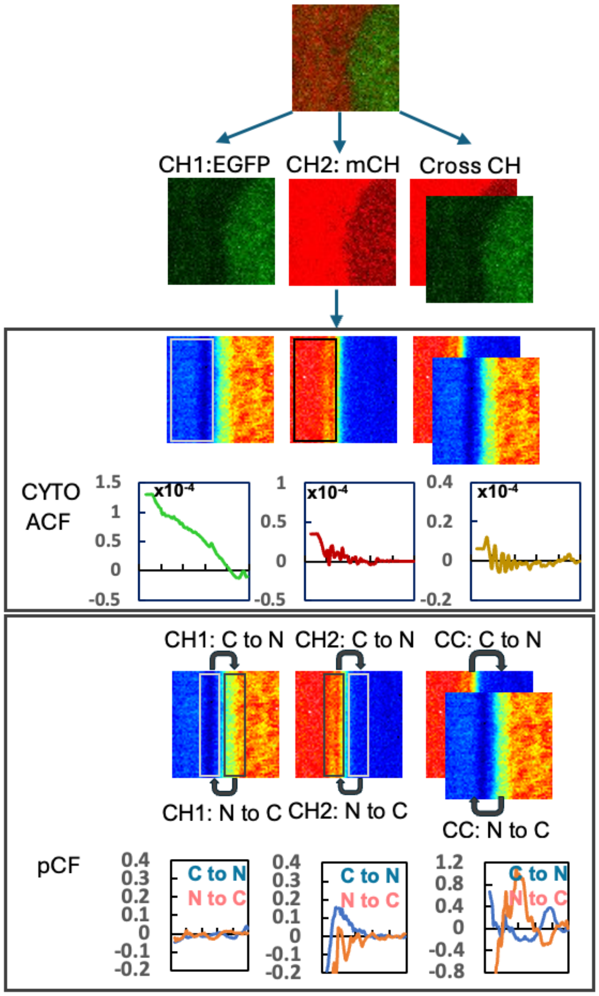
CH1:EGFP
CH2: mCH
Cross CH
CYTO
ACF
x10⁻⁴
x10⁻⁴
x10⁻⁴
CH1: C to N
CH2: C to N
CC: C to N
CH1: N to C
CH2: N to C
CC: N to C
pCF
C to N
N to C
C to N
N to C
C to N
N to C

#### Figure S6

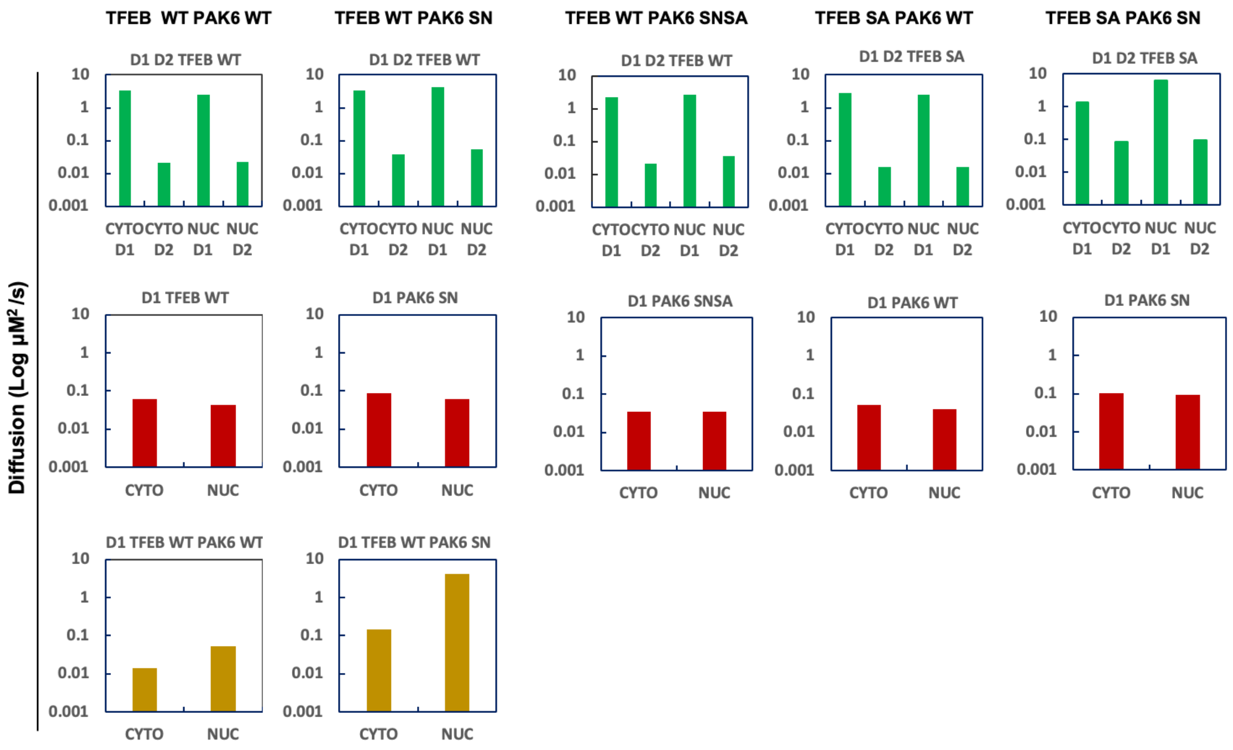
TFEB WT PAK6 WT
TFEB WT PAK6 SN
TFEB WT PAK6 SNSA
TFEB SA PAK6 WT
TFEB SA PAK6 SN
Diffusion (Log µM²/s)
D1 D2 TFEB WT
D1 D2 TFEB SA
10
1
0.1
0.01
0.001
CYTO D1
CYTO D2
NUC D1
NUC D2
D1 TFEB WT
D1 PAK6 SN
D1 PAK6 SNSA
D1 PAK6 WT
CYTO
NUC
D1 TFEB WT PAK6 WT
D1 TFEB WT PAK6 SN

### Figure S7

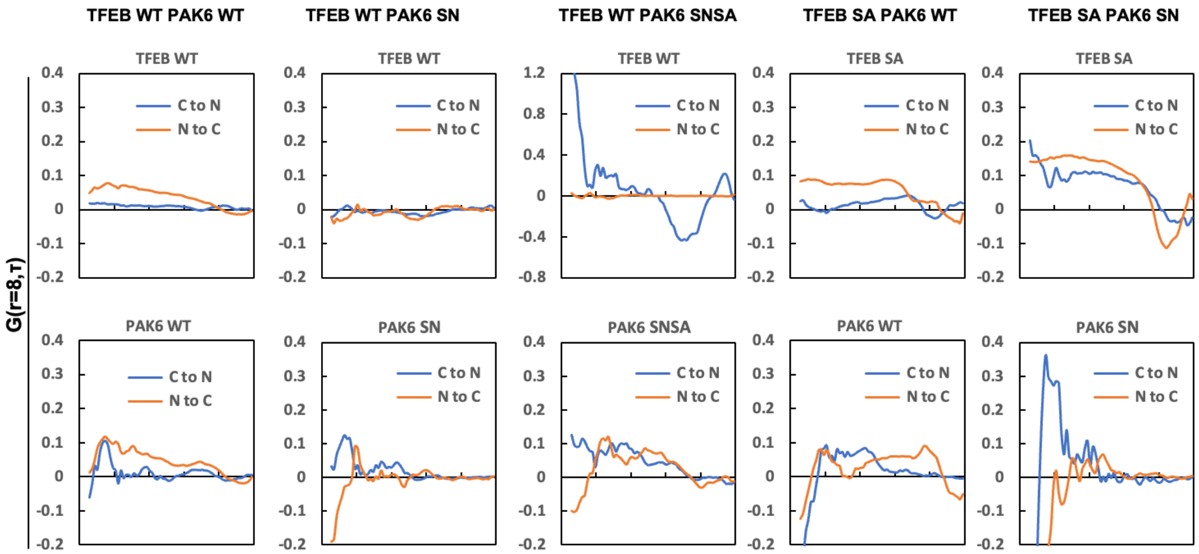


| | TFEB | | | | PAK6 | | Cross Channel (CC) | | Cross Channel (CC) | |
|---|---|---|---|---|---|---|---|---|---|---|
| | CYTO D1 | CYTO D2 | NUC D1 | NUC D2 | CYTO D1 | NUC D1 | CYTO D1 | NUC D1 | F CYTO | F NUC |
| TFEB WT PAK6 WT | 3.36±0.58 | 0.02±0.01 | 2.55+0.66 | 0.02±0.00 | 0.06±0.03 | 0.04±0.02 | 0.21±0.07 | 0.25±0.09 | 0.12±0.03 | 0.11±0.02 |
| TFEB WT PAK6 SN | 3.27±0.49 | 0.04±0.01 | 4.16±0.53 | 0.06±0.01 | 0.09±0.03 | 0.06±0.02 | 0.15±0.07 | 4.18±2.72 | 0.17±0.04 | 0.25±0.03 |
| TFEB WT PAK6 SNSA | 2.19±0.68 | 0.02±0.00 | 2.65±0.69 | 0.03±0.00 | 0.03±0.01 | 0.03±0.00 | | | | |
| TFEB SA PAK6 WT | 2.79±0.51 | 0.02±0.00 | 2.43±0.47 | 0.02±0.00 | 0.05±0.01 | 0.04±0.02 | | | | |
| TFEB SA PAK6 SN | 1.33±0.58 | 0.09±0.06 | 6.37±4.96 | 0.10±0.07 | 0.10±0.05 | 0.09±0.04 | | | | |

**Figure S8**

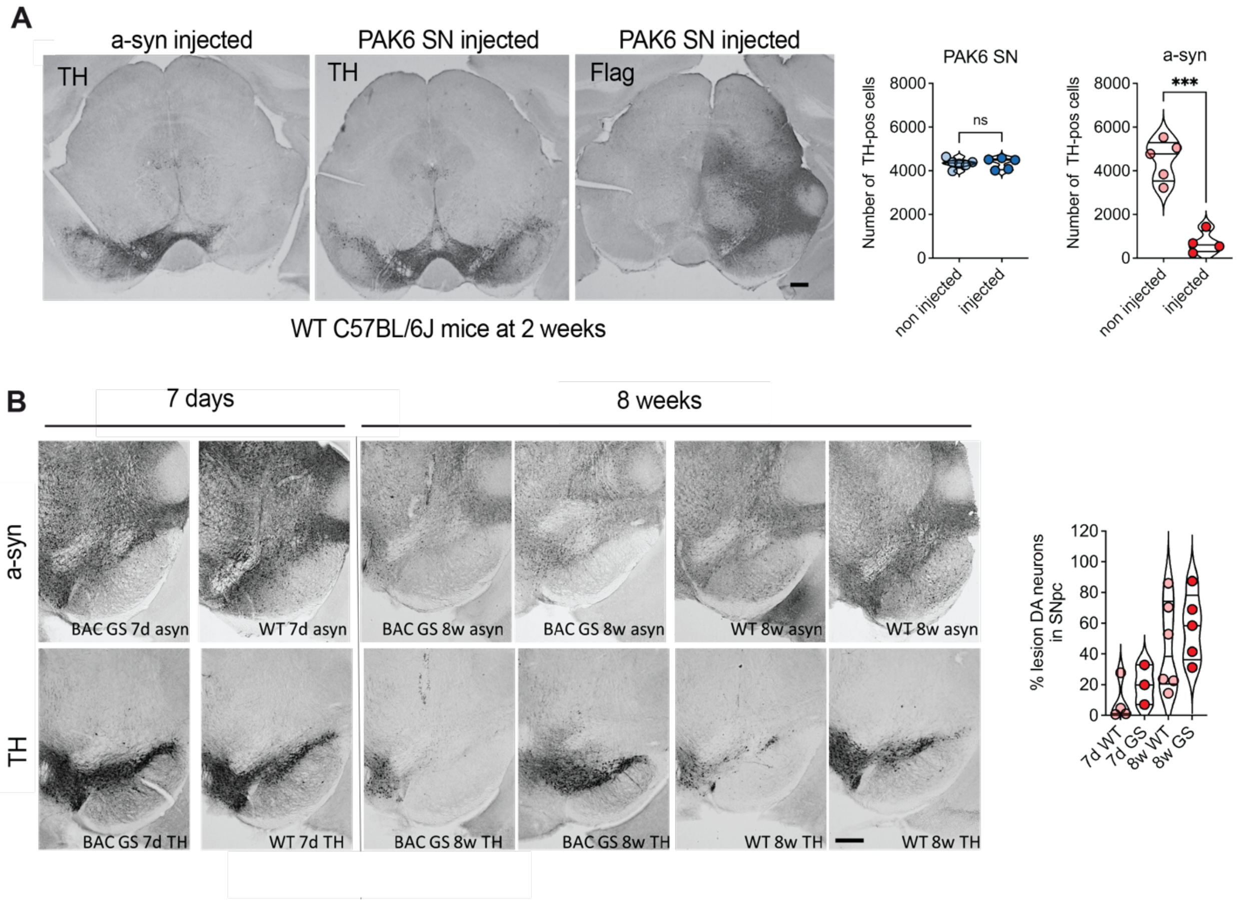
A
a-syn injected
PAK6 SN injected
PAK6 SN injected
TH
TH
Flag
WT C57BL/6J mice at 2 weeks
PAK6 SN
a-syn
Number of TH-pos cells
ns
***
non injected
injected
B
7 days
8 weeks
a-syn
TH
BAC GS 7d asyn
WT 7d asyn
BAC GS 8w asyn
WT 8w asyn
BAC GS 7d TH
WT 7d TH
BAC GS 8w TH
WT 8w TH
% lesion DA neurons in SNpc
7d WT
7d GS
8w WT
8w GS
